# Simultaneous therapeutic targeting of inflammation and virus ameliorates influenza pneumonia and protects from morbidity and mortality

**DOI:** 10.1101/2022.02.09.479486

**Authors:** Pratikshya Pandey, Zahrah Al Rumaih, Ma. Junaliah Tuazon Kels, Esther Ng, KC Rajendra, Roslyn Malley, Geeta Chaudhri, Gunasegaran Karupiah

## Abstract

Pneumonia is a severe complication caused by inflammation of the lungs following infection with seasonal and pandemic strains of influenza A virus (IAV) that can result in lung pathology, respiratory failure and death. There is currently no treatment available for severe disease and pneumonia caused by IAV. Antivirals are available, but they are far from satisfactory if treatment is not initiated within 48 hours of symptoms onset. Influenza complications and mortality are often associated with high viral load and excessive lung inflammatory cytokine response. Therefore, we simultaneously targeted IAV with the antiviral drug oseltamivir and inflammation with the anti-inflammatory drug etanercept, targeting TNF after the onset of clinical signs to treat IAV pneumonia effectively. The combined treatment effectively reduced lung viral load, lung pathology, morbidity and mortality during respiratory IAV infection in mice, contemporaneous with significant downregulation of the inflammatory cytokines TNF, IL-1β, IL-6, IL-12p40, chemokines CCL2, CCL5 and CXCL10 and dampened STAT3 activation. Consequently, combined therapy with oseltamivir and a STAT3 inhibitor also effectively reduced clinical disease and lung pathology. Combined treatment using either of the anti-inflammatory drugs and oseltamivir dampened an overlapping set of cytokines. Thus, combined therapy targeting a specific cytokine or cytokine signaling pathway plus an antiviral drug provides an effective treatment strategy for ameliorating IAV pneumonia. Effective treatment of IAV pneumonia required multiple doses of etanercept and a high dose of oseltamivir. This approach might apply to the treatment of pneumonia caused by severe acute respiratory syndrome coronavirus 2 (SARS-CoV-2).

**Significance Statement:** Antivirals against influenza A virus (IAV) are ineffective in treating pneumonia if administered 48 h after onset of disease symptoms. The host inflammatory response and tissue damage caused by IAV are responsible for lung pathology. We reasoned that targeting both virus and inflammation would be more effective in reducing lung pathology and pneumonia, morbidity and mortality. The simultaneous treatment with an anti-inflammatory drug targeting TNF or STAT3, combined with the anti-IAV antiviral drug, oseltamivir, significantly improved clinical disease, reduced lung viral load and pathology, and protected mice from severe pneumonia. The combined treatment suppressed multiple pro-inflammatory cytokines and cytokine signaling pathways. Thus, after the onset of disease symptoms, both virus and inflammation must be targeted to treat IAV pneumonia effectively.

## Introduction

Pneumonia is a serious complication caused by inflammation of the lungs following infection with seasonal and pandemic influenza viruses that can result in lung pathology, respiratory failure and death (1-4). There are currently no treatments for influenza pneumonia and antivirals against influenza A viruses (IAVs) are far from satisfactory if treatment is not initiated within 48 h of onset of disease symptoms (5). Most individuals do not seek medical attention within this timeframe (6). There is thus an urgent need to advance therapies that specifically treat severe IAV post-onset of symptoms.

An over-exuberant immune response associated with dysregulated inflammatory cytokine/chemokine production, known as a ‘cytokine storm’ (7), causes pneumonia, lung pathology and death (4). Late after onset of symptoms (>48 h), the damaging effects of inflammatory cytokines and virus-mediated cytopathic effects plus tissue necrosis together contribute to lung pathology, morbidity and mortality (8). Excessive early inflammatory cytokine/chemokine responses and leukocyte recruitment can be predictive of poor prognosis and poor clinical outcomes in IAV infections (1, 9, 10) and those inflammatory factors directly contribute to leukocyte recruitment into the lungs (8, 11, 12). We reasoned that the simultaneous targeting of both virus and inflammation would be an effective treatment strategy to ameliorate influenza pneumonia (13). Targeting inflammatory cytokines or cytokine-signaling molecules will reduce inflammation and diminish leukocyte infiltration into the lung.

Of the various cytokines implicated, tumor necrosis factor (TNF) is a crucial driver of inflammation in IAV-induced pneumonia (14-17). Viral infection triggers rapid TNF production by the innate immune system through activation of the nuclear factor-kappa B (NF-κB) signaling pathway (18). TNF exists in two distinct forms: soluble TNF (sTNF) and its precursor, transmembrane TNF (mTNF). Both forms of TNF can bind to their cognate receptors, TNF receptors type I and II (TNFRI and TNFRII) and mediate biological effects on various cell types, mainly through the activation of the NF-κB pathway (13, 19). Both TNFR also exist as soluble (sTNFR) and membrane-bound (mTNFR) forms. The binding of sTNFRII or mTNFRII to mTNF can also transmit signals into mTNF-bearing cells and dampen inflammation through a process known as ‘reverse signaling’ (20, 21).

Oseltamivir (22), an IAV neuraminidase (NA) inhibitor (NAI), is the most common anti-IAV drug currently used to treat individuals in high-risk groups (23). It is only effective if treatment is initiated within 48h of onset of disease symptoms (5, 24). Several clinical trials and meta-analyses have shown that oseltamivir is not very effective in reducing severe disease phenotypes, including hospitalization and pneumonia when treatment is commenced >48 h after the onset of symptoms (5).

We used a murine model of acute respiratory H1N1 IAV infection to investigate why oseltamivir is ineffective in reducing morbidity and mortality if treatment is initiated late after the onset of disease signs. We have found that oseltamivir can effectively reduce disease severity and mortality in IAV-infected mice even after the onset of disease signs only when inflammation is simultaneously also targeted. We used the anti-TNF drug etanercept, widely used to treat several inflammatory diseases (25). Etanercept alone reduced disease signs and lung pathology but did not protect mice from mortality. Similarly, treatment with only oseltamivir was effective in reducing viral load but animals died from severe lung pathology.

Dysregulated TNF production results in the dysregulation of an overlapping set of cytokines, chemokines and cytokine-signaling pathways, including the NF-κB and signal transducer and activator of transcription (STAT) 3 pathways (26, 27). We used an inhibitor of STAT3, S3I-201, as an alternative anti-inflammatory drug to reduce lung inflammation. Combined treatment with S3I-201 and oseltamivir reduced viral load, clinical illness, lung inflammation and pathology in IAV-infected mice. Notably, the combined treatment with oseltamivir and etanercept or S3I-201 dampened the expression levels of an overlapping set of inflammatory cytokines, chemokines, and phosphorylated STAT3 (pSTAT3) protein. Many of these factors have been implicated causing lung pathology during IAV infection.

Our findings not only help explain why NAIs are ineffective in treating severe disease and pneumonia caused by IAV infection if administered late after the onset of disease symptoms but also provide effective treatment strategies. Although excessive levels of several cytokines and chemokines have been implicated in the pathogenesis of influenza pneumonia and lung pathology, our results indicate that the targeting of just one inflammatory cytokine or a cytokine signaling pathway is sufficient to ameliorate lung pathology and protect mice from an otherwise lethal disease when treatment is combined with oseltamivir.

## Results

### Late after Disease Onset, High Viral Load Drives Inflammatory Cytokine Response Contributing to Disease Severity and Mortality in IAV-infected Mice

Excessive early cytokine responses can be predictive of poor prognosis and poor clinical outcomes (1, 9, 10). We hypothesized that the escalating viral load, late after disease onset, will potentially drive a higher inflammatory response. We wanted to first establish whether a high viral load, increased levels of inflammatory cytokines/chemokines or both contributed to morbidity and mortality of IAV infected mice.

Groups of C57BL/6 mice (n=5) were left uninfected (control) or infected with 1000, 2000 or 3000 PFU through the intranasal (i.n.) route and killed for ethical reasons when they had lost significant weight or were morbid, and the remaining animals were killed on day 12 post-infection (p.i.).

Disease severity, determined by the extent of weight loss and clinical scores (condition of hair coat, posture, breathing, lacrimation and nasal discharge, and activity and behavior as described elsewhere (26, 27) were generally significantly higher in mice infected with higher doses of IAV (Supplementary Fig. S1 *A* and *B*). Expectedly, viral load was higher in the lungs of mice infected with higher doses of IAV (Supplementary Fig. S1*C*). Regardless of the dose of virus inoculated, lung IAV load positively correlated with weight loss and clinical scores (Pearson’s coefficient r > 0.8) (Table 1). The survival rates were 60%, 20% and 40% for mice infected with 1000, 2000 and 3000 PFU, respectively (Supplementary Fig. S1*D)*. A total of 9 animals from the 1000 (n=2), 2000 (n=4) and 3000 (n=3) PFU infection groups that were severely morbid and succumbed early to infection (days 6-10 p.i.), had high lung viral load (>10^4^ PFU/g lung tissue), regardless of the dose of virus inoculation (Supplementary Fig. S1*E*). Except for two infected mice from the 3000 PFU infection group, all remaining animals killed on day 12 p.i. had a low lung viral load. Thus, high lung viral load was associated with increased disease severity and mortality in IAV-infected mice (Supplementary Fig. S1*E)*.

**Table 1.** Correlation coefficient between disease severity determinants and lung viral load or levels of pro-inflammatory mediators in IAV-infected mice. Age-matched groups of WT mice (n = 5) were infected with 1000, 2000, or 3000 PFU IAV i.n. Weight loss and clinical scores were assessed until day 12 p.i., when all the animals were killed. Lungs were collected for measuring viral load and mRNA transcripts for pro-inflammatory cytokines and chemokines. Data shown in Fig. S1 is presented here as Pearson correlation coefficient (r) between any of the two determinants of disease severity, namely correlation between weight loss and lung viral load or mRNA levels of pro-inflammatory mediators, and correlation between clinical scores and lung viral load or mRNA levels of pro-inflammatory mediators.

| Disease severity determinants | Viral load | Cytokines and chemokines |  |  |  |  |  |
| --- | --- | --- | --- | --- | --- | --- | --- |
|  |  | TNF | IL-6 | IL-12p40 | CCL2 | CCL5 | CXCL10 |
| Weight loss | 0.84 | 0.79 | 0.72 | 0.62 | 0.57 | 0.68 | 0.72 |
| Clinical score | 0.82 | 0.74 | 0.72 | 0.57 | 0.52 | 0.68 | 0.74 |

The levels of mRNA transcripts for TNF, IL-6, IL-12p40, CCL2, CCL5, and CXCL10 positively correlated (Pearson’s coefficient r > 0.5) (Table 1) with weight loss and clinical scores (Supplementary Fig. S1 *F-K*). In particular, increasing the virus inoculum dose increased IL-6, CCL2, CCL5, and CXCL10 mRNA levels, and animals with high levels of cytokines/chemokines were more likely to become moribund and die compared (Supplementary Fig. S1 L-Q). All 9 mice that succumbed to infection expressed high levels of cytokines and chemokines. Importantly, two infected mice that survived despite having high viral titers had lower cytokine/chemokine levels than those which succumbed. Thus, high viral load in parallel with high cytokine/chemokine transcript levels is associated with increased morbidity and mortality of IAV infected mice.

### Etanercept Treatment of WT Mice Improves Clinical Disease and Reduces Lung Immunopathology without Affecting IAV Load but TNF Deficiency Exacerbates Lung Pathology

Etanercept mediates anti-inflammatory effects by neutralizing sTNF and triggering reverse signaling via mTNF (28, 29). We first used WT mice and a triple mutant (TM) strain, which expresses only the non-cleavable mTNF but not sTNF, TNFRI and TNFRII, to establish that etanercept can reduce lung inflammation during an IAV infection. TM mice lack endogenous TNF signaling like TNF^-/-^ mice but respond to exogenous TNFR such as etanercept. WT and TM mice infected with 3000 PFU IAV i.n. were treated with 2.5 mg/kg etanercept or vehicle (PBS) intraperitoneally (i.p.) on days 1, 3 and 4 p.i. Animals were killed on day 5 p.i.

Compared with WT mice, mock-treated TM mice developed more severe disease as evident by higher losses in body weights and increased clinical scores and lung histopathological scores, evaluate pulmonary inflammatory cell infiltrate, oedema and bronchial epithelial cell loss as markers of inflammation and lung damage (Fig. 1 *A-E*). We generated lung histopathological scores from the microscopic examination of hematoxylin and eosin (H&E)-stained lung sections described elsewhere (26, 27). Treatment with etanercept significantly reduced weight loss, clinical scores and histopathological scores in both strains of mice (Fig. 1 *A-E*) but did not affect viral load (Fig. 1*F*). Microscopic examination of lung histological sections revealed dramatic reductions in parenchymal edema and inflammatory cell infiltration by etanercept treatment, consistent with reduced weight loss, clinical scores and histopathological scores (Fig. 1*G*).

**Fig. 1.**
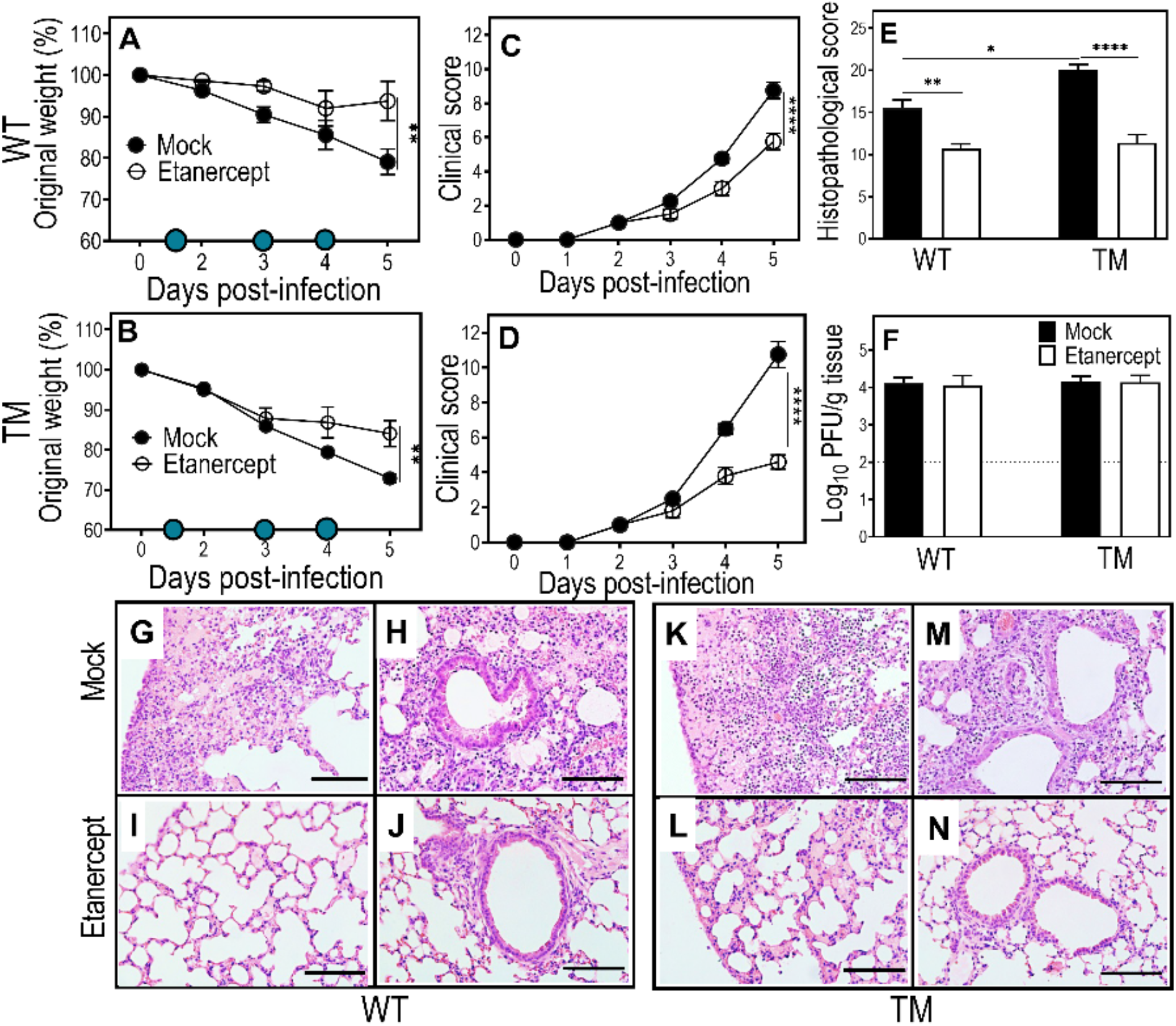
Etanercept treatment reduces weight loss, clinical scores and lung pathology but not viral load in IAV-infected WT and TM mice. Age-matched groups of female WT and TM mice (n = 4 or 5) were infected with 3000 PFU IAV i.n. Animals were treated with 2.5 mg/kg etanercept or diluent (mock) on days 1, 3 and 4 p.i. as indicated in panels A and B where filled blue circle symbols indicate etanercept treatment days. Animals were killed on day 5 p.i. and lungs collected for analyses. Weight loss (A and B) and clinical scores (C and D) were analyzed using two-way ANOVA with Sidak’s post-tests and expressed as means ± SEM. Viral load data (E) were log-transformed, analyzed using ordinary one-way ANOVA test with Fisher’s least significant difference (LSD) tests and expressed as means ± SEM. Histopathological scores (F), based on microscopic examination of lung histology H&E sections (G), were examined using bright field microscope on all fields at 400x magnification. Histopathological scores were analyzed by ordinary one-way ANOVA test with Tukey’s post-tests and expressed as means ± SEM. Lung histology sections (G-N) show reductions in edema, leukocyte infiltration, and damage to alveolar septa in lungs of IAV-infected WT and TM mice by etanercept treatment. *, p < 0.05; **, p < 0.01 and ***, p <0.001. Broken line in panel E corresponds to the limit of virus detection. Bars in panel G-N correspond to 100 µ m. TM, triple mutant mice that express mTNF but not sTNF, TNFRI or TNFRII; WT, wild-type mice. Data shown are from a single experiment.

We tested the response of TNF^-/-^ mice to IAV infection as we wanted to use them as controls in experiments with etanercept and oseltamivir combined treatment. We found that IAV-infected TNF^-/-^ mice had significantly higher clinical scores than WT mice, evident from days 4-6 p.i. although both strains had comparable body weight losses from days 2-6 p.i. (Supplementary Fig. S2 *A* and *B*). The lung viral load was high (>10^6^ PFU) and comparable in both strains of mice (Supplementary, Fig. S2*C*). TNF^-/-^ mice had significantly higher histopathological scores (Supplementary Fig. S2*D*) due to the more pronounced lung pathological changes compared to WT mice (Supplementary Fig. S2 *E-P*), consistent with a previous report (30).

### One Dose of Etanercept Combined with a Standard Dose of Oseltamivir (40 mg/kg) Daily Treatment Reduces Morbidity and Lung Pathology but has no Effect on Viral Load

Etanercept treatment beginning at day 1 p.i. with IAV improved clinical disease and lung pathology (Fig. 1). However, most patients seek medical attention late after the onset of symptoms (6). Furthermore, the cytopathic effects of exponentially increasing viral load also contribute to lung injury at that stage of the disease, but etanercept does not affect lung viral load. We reasoned that combined treatment with etanercept and oseltamivir simultaneously would be necessary to minimize disease severity effectively.

An oral dose of 150 mg oseltamivir twice daily is well-tolerated in adult humans (31) and is equivalent to a dose of 20 mg/kg in mice administered twice daily (32). Mice infected with 3000 PFU IAV i.n. were given an oral dose of oseltamivir at 20 mg/kg (twice daily; total 40 mg/kg/day), one dose of etanercept, or both drugs (combined) on day 3 p.i. after onset of disease signs. Additional doses of oseltamivir were given on days 4 and 5 p.i. All mice were killed on day 6 p.i. Since the clinical course of IAV disease is short in mice, it leaves a very narrow window period for treatment. We commenced treatment from day 3 p.i, so that animals receive appropriate treatment(s) for at least 2 days before they succumb to infection.

Oseltamivir or combined treatment significantly reduced weight loss, clinical scores and lung histopathological scores (Supplementary Fig. S3 *A-C*) but did not affect viral load in the lung (Supplementary Fig. S3*D*). A single dose of etanercept also significantly reduced clinical scores but had no impact on weight loss or histopathological scores (Supplementary Fig. S3 *E-I*).

Oseltamivir or etanercept significantly reduced the levels of mRNA transcripts for TNF, IL-6, IL-12p40, CCL2, and CCL5 (Supplementary Fig. S4 *A* and *C-F*). In addition to these cytokines/chemokines, the combined treatment significantly downregulated levels of expression of IL-1β and CXCL10 (Supplementary Fig. S4 *B* and *G)*. mTNF levels were significantly reduced by all three treatment regimens (Supplementary Fig. S4*H)*. IAV infection minimally activated NF-κB p65, but levels were significantly reduced by oseltamivir or combined treatment (Supplementary Fig. S4*I)*. On the other hand, STAT3, another critical transcription factor implicated in the host inflammatory response, was highly activated by IAV infection. All three treatment regimens reduced IAV-induced pSTAT3 levels, however, the reduction was modest with etanercept or combined treatments but was significant with oseltamivir treatment alone (Supplementary Fig. S4*J)*. In terms of lung levels of pSTAT3, we observed high variability between animals in each of the three treatment groups.

### Combined Daily Treatment with Etanercept and High Dose Oseltamivir Reduces Morbidity, Lung Viral Load and Pathology in IAV-infected Mice

A single dose of etanercept and 20 mg/kg oseltamivir (administered twice daily) were ineffective in reducing weight loss, viral load and histopathological scores. Chronic inflammatory diseases like rheumatoid arthritis are treated with etanercept once or twice a week (25). However, levels of TNF produced during an acute viral infection is likely to be different to those during a chronic inflammatory condition. Therefore, we have administered etanercept and 150 mg/kg oseltamivir once daily in subsequent experiments, unless indicated otherwise.

Groups of IAV-infected WT, TM and TNF^-/-^ mice were treated with oseltamivir, etanercept or both drugs after onset of disease signs on day 3 p.i., treatment continued on days 4 and 5 p.i. once daily and animals killed at day 6 p.i. TNF^-/-^ mice were used as negative controls as they do not respond to etanercept treatment (33) (33). TM mice lack endogenous TNF signaling but respond to etanercept treatment (Fig. 1).

Mock-, etanercept-or oseltamivir-treated WT mice continued to lose weight until day 6 p.i. when animals were killed for various analyses. In contrast, WT mice given the combined treatment stopped losing weight just one day after initiation of treatment and by day 6 p.i., body weight loss in this group was significantly lower than the other groups (Fig. 2*A*). All treatment regimens significantly lowered clinical scores from day 4 p.i., but the most significant reduction was in the combined treatment group (Fig. 2*B*). Lung viral load was similar in mock, and etanercept treated groups but significantly reduced by oseltamivir or combined treatment compared with the mock-treated group (Fig. 2*C*). Notably, once-daily treatment with150 mg/kg oseltamivir alone effectively reduced viral load late after the onset of disease signs. There were significant reductions in lung histopathological scores in etanercept-but not oseltamivir-treated animals compared to mock-treated mice, but the combined treatment showed the most considerable reduction (Fig. 2*D*).

**Fig. 2.**
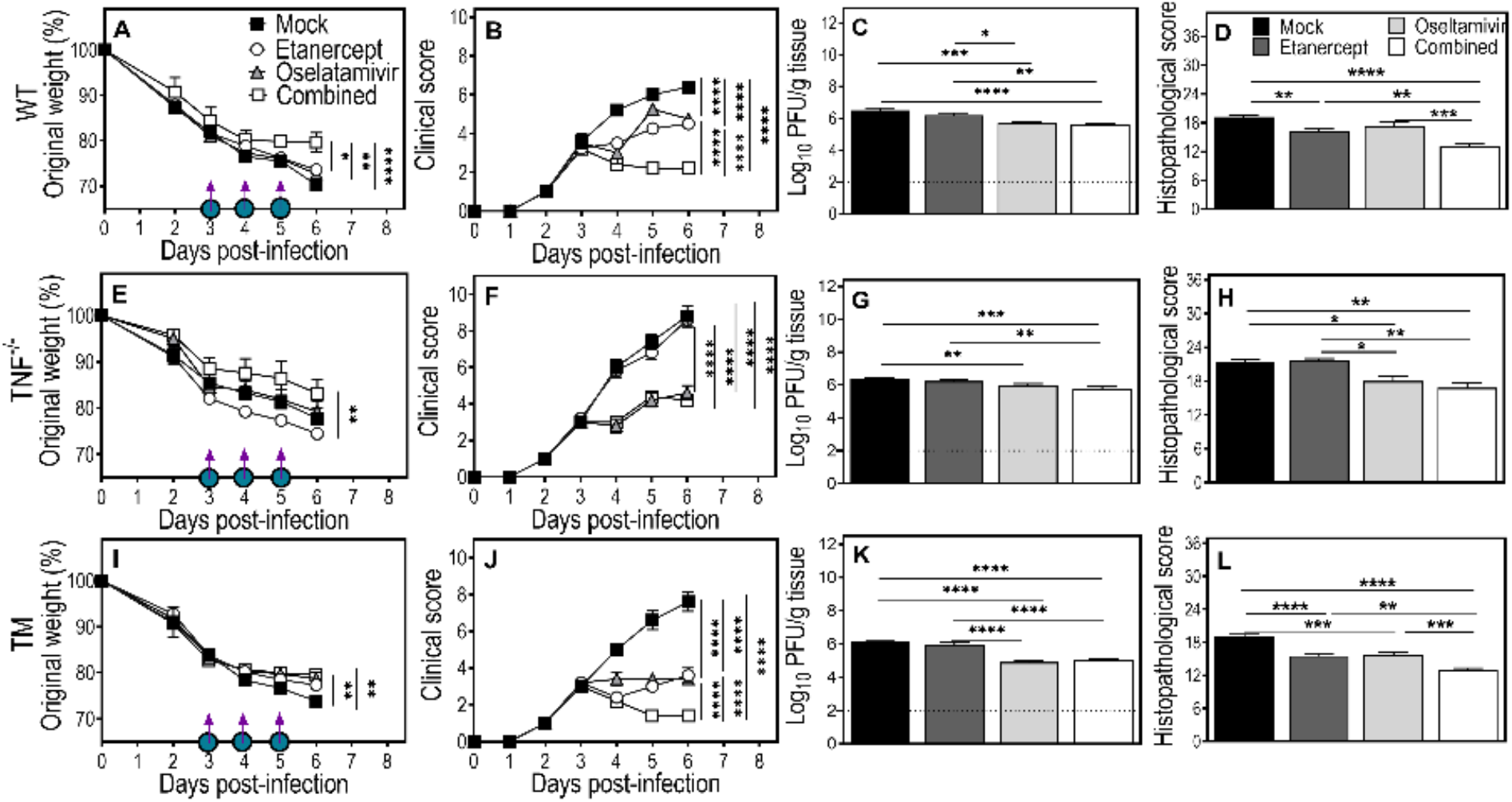
Combined treatment with etanercept and oseltamivir reduces clinical scores, lung viral load and pathology in IAV-infected WT, TNF^-/-^ and TM mice. Age-matched groups of WT, TNF^-/-^ and TM (n = 4 or 5) female mice were infected with 3000 PFU IAV i.n., treated with oseltamivir (150 mg/kg), etanercept (2.5 mg/kg) or a combination (combined) on days 3, 4 and 5 p.i., as indicated in panels A, E and I where a purple arrow and a filled blue circle symbols indicate oseltamivir and etanercept treatment days, respectively. Animals were monitored for weight loss (A, E and I) and clinical scores (B, F and J) until day 6 p.i., when all animals were killed and lung tissue collected for various analyses. Data were analyzed by two-way ANOVA with Sidak’s post-tests and expressed as means ± SEM. Viral load (C, G and K) data was log-transformed, analyzed using ordinary one-way ANOVA test followed by Fisher’s LSD tests and expressed as means ± SEM. Histopathological scores (D, H and L) were derived from microscopic examination of lung histology H&E sections (presented in Fig. 3), analyzed using ordinary one-way ANOVA test followed by Tukey’s multiple comparisons tests and expressed as means ± SEM. *, p < 0.05; **, p < 0.01; ***, p <0.001 and ****, p <0.0001. Broken lines in panels C, G and K correspond to the limit of virus detection. Data shown are from a single experiment.

In IAV-infected TNF^-/-^ mice, none of the treatment regimens had any significant effects in reducing weight loss (Fig. 2*E*). Mice given oseltamivir or the combined treatment had significantly lower clinical scores than etanercept-or mock-treated animals (Fig. 2*F*). Lung viral load and histopathological scores were also reduced by oseltamivir or combined treatment but not by etanercept-or mock treatment (Fig. 2 *G* and *H*). The reduced viral load likely contributed to the lower clinical and histopathological scores in TNF^-/-^ mice.

TM mice exhibited apparent beneficial effects from the single or combined treatment regimens, similar to observations made in WT mice (Fig. 2 *I-L*). On day 6 p.i., etanercept treatment reduced weight loss compared to mock treatment, but the effect was more pronounced in oseltamivir or combined treatment groups (Fig. 2*I*). All treatment regimens significantly reduced clinical scores, with the combined treatment having the most prominent effect (Fig. 2*J*). Oseltamivir or combined treatment, but not etanercept, reduced lung viral load (Fig. 2*K*). Lung histopathological scores were significantly reduced by oseltamivir or etanercept monotherapy, but the combined treatment reduced it to a substantially greater extent (Fig. 2*L*).

Microscopic examination of lung histological sections revealed dramatic reductions in parenchymal edema and damage to bronchial and alveolar walls in WT mice given the combined treatment compared to the other treatment groups (Fig. 3 *A-H*). Focal leukocyte infiltration was most abundant in lungs of WT mice given mock-treatment compared with oseltamivir-or etanercept-treated groups, but it was only moderate in mice given the combined therapy (Fig. 3 *A-H*). In TNF^-/-^ mice, IAV-induced lung edema and inflammatory cell recruitment, were reduced by oseltamivir or combined treatment, but not by etanercept (Fig. 3 *I-P*). However, the extent of improvements was less compared to WT mice. All treatment regimens ameliorated edema and inflammatory cell infiltration in TM mice, except for mock treatment, but the degree of improvement was highest in the combined-treated group (Fig. 3 Q*-X*).

**Fig. 3.**
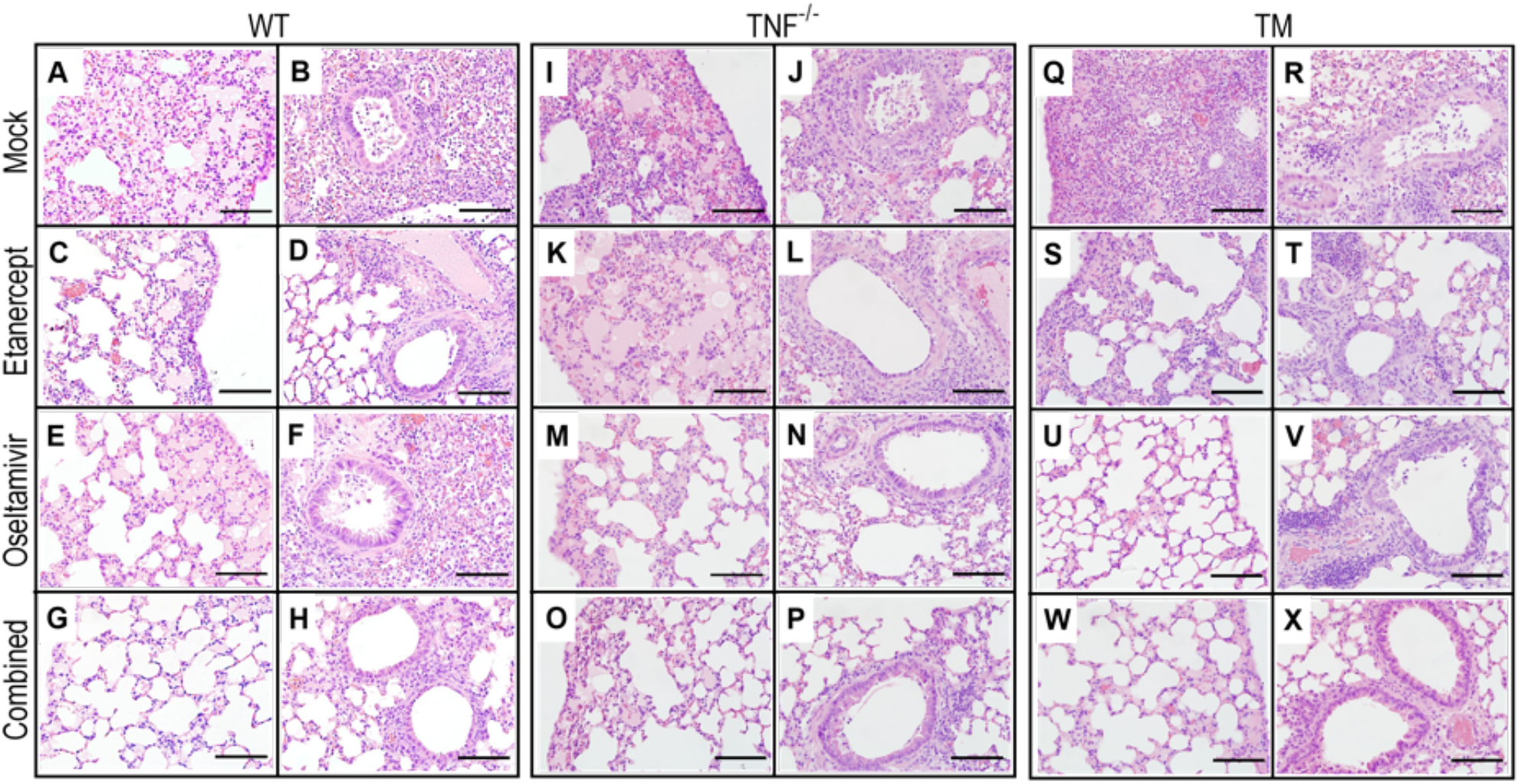
Combined daily treatment with etanercept and high dose oseltamivir reduces IAV-infection induced lung pathology to a greater extent in WT and TM mice than TNF^-/-^ mice. Lung tissue sections were obtained from mice that were infected and treated as described in Fig. 2. Briefly, groups of WT, TNF^-/-^ and TM mice (n = 4 or 5) were infected with 3000 PFU IAV i.n. and then treated with oseltamivir or etanercept or both drugs combined on days 3, 4 and 5 p.i. Animals were killed on day 6 p.i., lungs were collected, fixed in 10% neutral buffered formalin, processed, embedded in paraffin blocks, sectioned, stained with H&E and examined using bright field microscope on all fields at 400x magnification. Bars in panels A-X correspond to 100 μm. Data shown are from a single experiment.

To obtain an insight into the mechanisms through which treatment with etanercept combined with a higher dose of oseltamivir, both administered daily (Fig. 2 and 3), we focused on the effects of treatment on inflammatory cytokine and chemokine mRNA transcript levels using lung tissue samples from the experiment described in Fig. 2.

Compared to mock treatment, etanercept or oseltamivir reduced mRNA transcripts for TNF, IL-1β, and IL-12p40 (Fig. 4 *A, B* and *C*), whereas the combined treatment was more effective in decreasing the levels of those cytokines as well as IL-6 and the chemokines CCL2, CCL5, and CXCL10 (Fig. 4 *D-G*). Levels of mTNF were decreased marginally by etanercept but significantly by oseltamivir or the combined treatment (Fig. 4*H*). All treatment regimens reduced levels of activated phosphorylated (p) NF-κB p65 (pNF-κB p65) protein, but only oseltamivir treatment was significant (Fig. 4*I*). However, protein levels of pSTAT3 were reduced only by oseltamivir or the combined treatment but not etanercept (Fig. 4*J*).

**Fig. 4.**
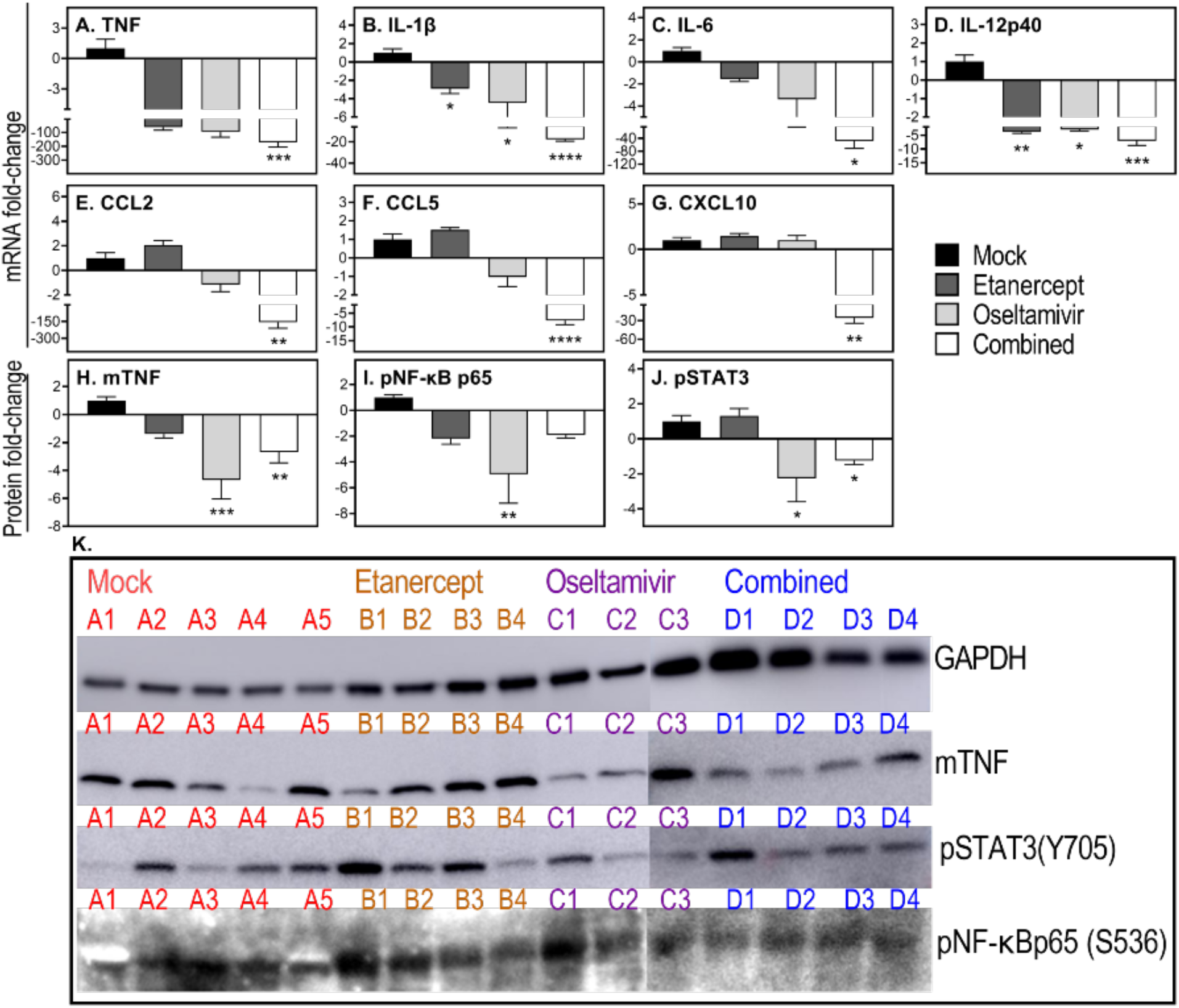
Combined daily treatment with etanercept and high dose oseltamivir reduces expression of inflammatory cytokines and chemokines and activation of STAT3. Lung tissues were obtained from WT mice that were infected and treated as described in Fig. 2. Briefly, WT (n = 4 or 5) mice were infected with 3000 PFU IAV i.n. and treated with oseltamivir (150 mg/kg) or etanercept or combined treatment on days 3, 4, and 5 p.i. Animals were killed on day 6 p.i. and lungs collected for quantifying levels of expression of selected cytokines and chemokines using qPCR (A-G). Protein levels of mTNF (H), pNF-κB p65 (I), and pSTAT3 (J) were detected by western blotting (K) and quantified with the ImageJ software. For (K), samples were run on 2 separate gels, i.e., gel 1, A1-C2; gel 2, C3-D4. Data were analyzed by one-way ANOVA with Holm-Sidak’s multiple comparisons tests and expressed as mean fold-change relative to the mock treated group ± SEM. *, p < 0.05; **, p < 0.01; ***, p <0.001 and ****, p <0.0001. Data shown are from a single experiment.

Taken together, the effectiveness of the combined treatment regimen in reducing weight loss, clinical disease, lung pathology and viral load was associated with significant reductions in TNF, IL-1β, IL-6, IL-12p40, CCL2, CCL5 and CXCL10 and to some extent pSTAT3. It was evident that the higher dose of oseltamivir resulted in significant reductions in viral load and levels of IL-1β more effectively (Fig. 4) compared to treatment with a lower dose oseltamivir (Supplementary Fig. S4).

### Combined Daily Treatment with Etanercept and High Dose Oseltamivir Protects Mice from Lethal IAV infection

We determined whether the higher dose of oseltamivir administration in the combined treatment regimen afforded protection against influenza pneumonia, and lethal disease. IAV-infected mice were treated with high dose oseltamivir, etanercept or both drugs after the onset of disease signs at day 3 p.i. once daily for up to 20 days. Animals were monitored for morbidity and mortality until day 21 p.i. when all surviving animals were killed. TNF^-/-^ mice, which do not respond to etanercept treatment (Fig. 2) were infected and treated similarly for use as controls.

Both WT and TNF^-/-^ mice infected with IAV continued to lose weight from day 2 p.i. and exhibited clinical signs of disease from day 3 p.i. (Fig. 5 *A-D*). All mock-treated WT mice succumbed to infection and were killed for ethical reasons by day 6 p.i. Etanercept treatment prolonged survival by 1 day in 4 of 5 (80%) IAV-infected WT mice (Fig. 5*E*). Oseltamivir treatment alone protected 1 of 5 (20%) mice from lethal IAV infection while the remaining animals succumbed on days 6 and 7 p.i. Mice given the combined treatment had the highest survival rate wherein one mouse succumbed to the infection on day 11 p.i. but 80% of animals survived until day 21 when they were killed (Fig. 5>*E*). Mock-treated TNF^-/-^ mice succumbed to the infection on days 5 or 6 p.i. (Fig. 5*F*) and there were no beneficial effects of etanercept, oseltamivir or the combined therapies as all IAV-infected animals succumbed by day 7 p.i.

**Fig. 5.**
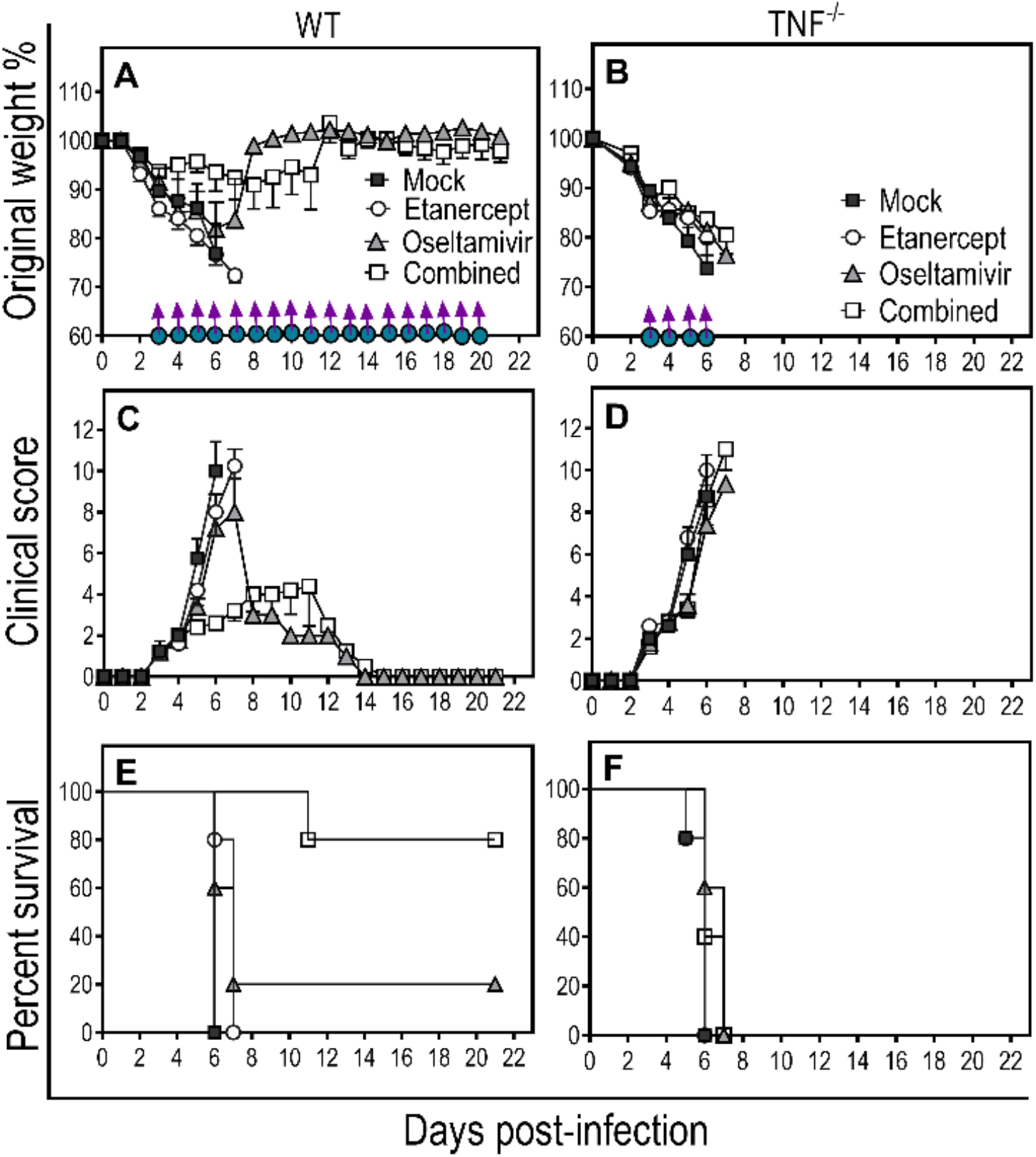
Combined daily treatment with etanercept and high dose oseltamivir reduces weight loss, and clinical scores and improves survival rate of WT but not TNF^-/-^ mice infected with IAV. Age-matched groups of WT and TNF^-/-^ (n = 4 or 5) mice were infected with 3000 PFU IAV i.n. Animals were treated with oseltamivir or etanercept or both drugs (combined) from day 3 p.i and treatment continued until day 20 p.i., as indicated in panels A and B, where a purple arrow and a filled blue circle symbols indicate oseltamivir and etanercept treatment days, respectively. Animals were monitored daily for weight loss (A and B), clinical scores (C and D) and survival (E and F). All mock treated WT mice died by day 6 p.i. whereas etanercept treated animals succumbed between days 6-7 (E). Four of 5 oseltamivir treated WT mice succumbed between days 6-7 p.i. whereas 1 animal was alive at day 21 p.i. The combined treatment resulted in 80% of WT mice surviving at day 21 (E). The median survival time for mice treated with etanercept or oseltamivir was 7 days, compared with a median survival of 6 days for mock-treated mice (log-rank test, mock vs etanercept p = 0.0237; mock vs oseltamivir, p = 0.0736). Survival for combined treated animals was greater than 50% (i.e., 80%) at the last time point (day 21 p.i.), hence the median survival time was >21 days (log-rank test, p = 0.0047 relative to mock-treated mice). Combined treatment significantly increased median survival compared to etanercept (p = 0.0035) or oseltamivir (p = 0.0358) treatments. Mock-treated TNF^-/-^ mice succumbed to infection on days 5-6 p.i. (F) and there were no beneficial effects of the different treatment regimens as all IAV-infected mice succumbed by day 7 p.i. Weight loss (A and B) and clinical scores (C and D) data were analyzed using two-way ANOVA with Sidak’s post-tests and expressed as means ± SEM. Survival data (E) were analyzed by Log-rank (Mantel-Cox) test. Data shown are from a single experiment.

Data presented in the preceding section indicated that viral load in WT mice was significantly reduced by oseltamivir alone or combined treatment regimens (Fig. 2*C*). However, a greater proportion of mice in the combined treatment group survived (Fig. 5*E*). It was clear that although high dose oseltamivir was effective in reducing viral load late after the onset of disease signs, it alone was insufficient to protect against influenza pneumonia, morbidity or mortality. Thus, both virus and inflammation must be targeted simultaneously to afford protection against influenza pneumonia.

### STAT3 Inhibitor in Combination with Oseltamivir Reduces Lung Viral Load and Improves Lung Pathology and Morbidity Associated with Severe IAV Infection

The STAT3 pathway is downstream of the NF-κB pathway, and dysregulated TNF levels cause hyperactivation of STAT3, which also correlated with severe pneumonia during respiratory ectromelia virus (ECTV) infection (26, 27). ECTV causes mousepox, a surrogate mouse model for smallpox caused by the variola virus in humans (34). Treatment of ECTV-infected mice with a selective STAT3 inhibitor, SI-301, significantly reduced lung pathology, but it was insufficient to protect the animals (26). However, inhibition of STAT3 combined with an antiviral drug, cidofovir, significantly reduced clinical disease and viral load and ameliorated lung pathology (33). The reduced morbidity was associated with reductions in inflammatory cytokine/chemokine gene/protein expression. We hypothesized that a similar approach of simultaneous targeting of virus and the STAT3 signaling pathway might also be effective in the IAV pneumonia model.

Although the standard dose of oseltamivir (40 mg/kg) was ineffective in reducing viral load when combined with etanercept (Supplementary Fig. S3*D*), we first investigated whether it would be effective in reducing morbidity when combined with S3I-201. Mice infected with IAV were given 20 mg/kg oseltamivir orally twice daily, 5mg/kg S3I-201 via the i.p. route or a combination of both drugs beginning on day 3 after the onset of disease signs, and treatment continued on days 4 and 5 p.i. All animals were killed on day 6 p.i.

The combined treatment significantly reduced weight loss, clinical scores, and lung histopathological scores but did not affect lung viral load (Supplementary Fig. S5 *A-I*). SI-301 had no effect on weight loss or viral load but reduced the clinical and histopathological scores. Mechanistically, oseltamivir reduced mRNA levels for IL-1β and IL-12p40 and protein levels of mTNF and pNF-κB (Supplementary Fig. S6 *B, D, H and I)*. S3I-201 or combined treatment significantly reduced mRNA transcripts for IL-1β, IL-6 and IL-12p40 (Supplementary Fig. S6 *B-D)*. In addition, the combined treatment also significantly reduced TNF and CCL2 mRNA and proteins levels of pNF-κB p65 and pSTAT3 (Supplementary Fig. S6 *A, E, I* and *J*). None of the treatment regimens had any significant impact on CCL5 and CXCL10 mRNA transcript (Supplementary Fig. S6 *F* and *G)*.

We next investigated whether the combined treatment regimen with S3I-201 and high dose oseltamivir would be more effective in reducing morbidity. Groups of IAV infected mice were treated with 150 mg/kg oseltamivir, 5mg/kg S3I-201 or both drugs (combined) on days 3 and 4 p.i. after the onset of disease signs. Since one of the mock-treated animals was severely moribund, all mice were killed on day 5 p.i. for ethical reasons.

High dose oseltamivir did not affect weight loss and modestly reduced lung histopathological scores but had a significant impact in reducing clinical scores and lung viral load (Fig. 6 *A-D*). S3I-201 or combined treatment significantly reduced weight loss, clinical scores and lung histopathological scores, while the latter also reduced the lung viral load and was by far the most effective in reducing all parameters measured (Fig. 6 *A-D*). Compared to other treatment groups, the combined treatment diminished lung inflammation and damage, evident by reduced edema, cellular infiltration, and bronchial epithelial necrosis (Fig. 6 *E-L*).

**Fig. 6.**
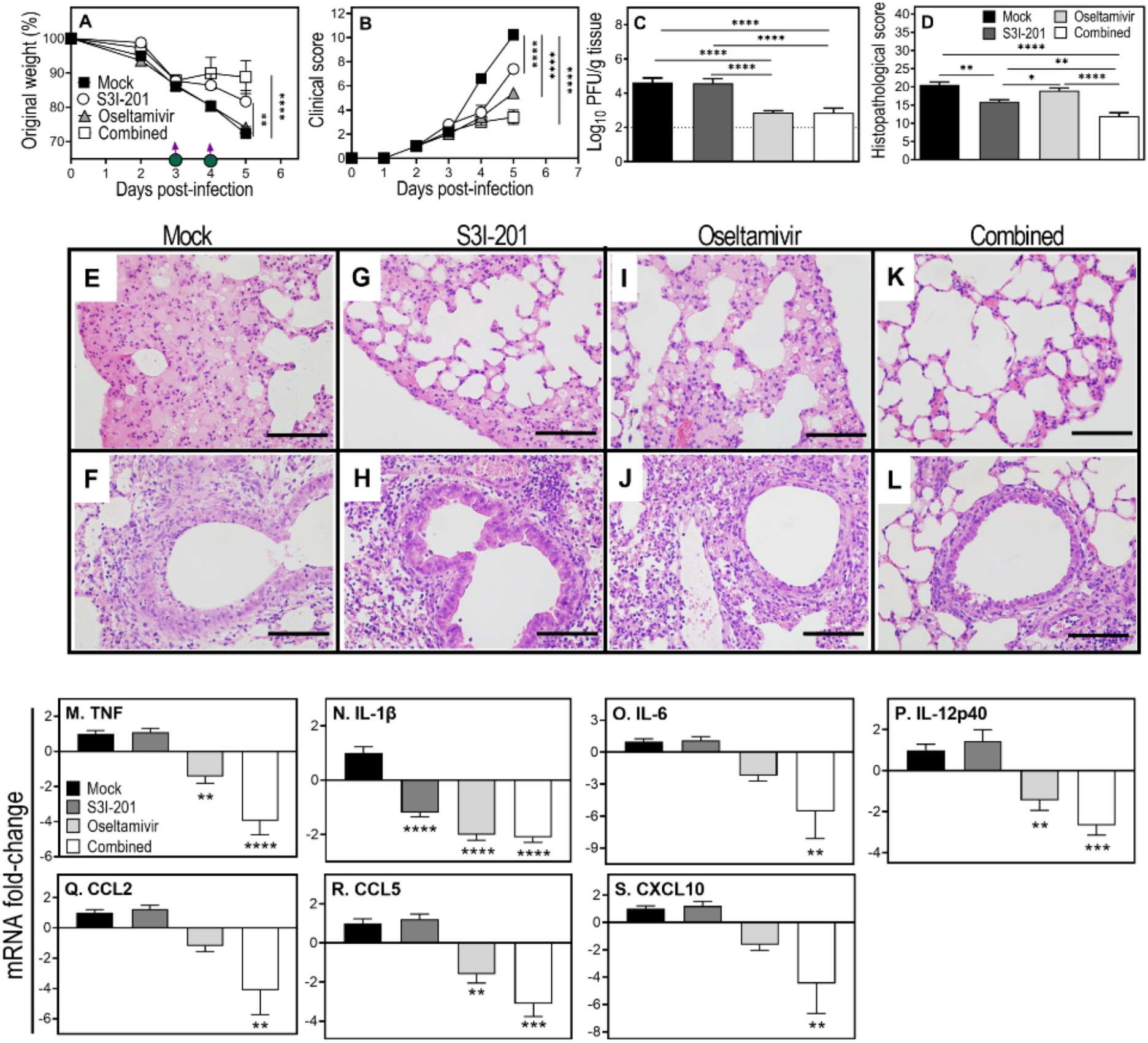
Combined treatment with high dose oseltamivir and S3I-201 reduces clinical scores, lung viral load, pathology and downregulates mRNA transcripts for pro-inflammatory cytokines and chemokines in IAV-infected mice. Age-matched groups (n = 5) of female WT mice were infected i.n. with 3000 PFU IAV and treated with S3I-201 (5 mg/kg), oseltamivir (150 mg/kg), or both drugs (combined) on days 3 and 4 p.i., as indicated in panel A, where a purple arrow and a filled green circle symbols indicate oseltamivir and S3I-201 treatment days, respectively. For ethical reasons, animals were killed on day 5 p.i. and lung tissue collected for various analyses. Weight loss (A) and clinical scores (B) were monitored until day 5 p.i. Viral load (C) data was log-transformed and analyzed using ordinary one-way ANOVA with Fisher’s LSD post-tests. Histopathological scores (D) were derived from microscopic examination of the lung histology H&E sections using bright field microscope on all fields at 400x magnification, presented in panel E-L. Data are expressed as means ± SEM and were analyzed using two-way ANOVA (A and B) with Tukey’s multiple comparisons tests (A) and Dunnett’s multiple comparisons tests (B) or ordinary one-way ANOVA followed by Tukey’s multiple comparisons tests (D). mRNA transcript analysis was performed in the separate experiment using the similar treatment strategy as described above. Gene expression levels of the indicated cytokines and chemokines were quantified using qPCR (M-S). Data are expressed as mean fold-change relative to the mock treated group ± SEM and were analyzed using one-way ANOVA with Holm-Sidak’s multiple comparisons tests. *, p < 0.05; **, p < 0.01; ***, p <0.001 and ****, p <0.0001. Broken line in panel C corresponds to the limit of virus detection. Bars in panels E-L correspond to 100 μm. Data shown are from a single experiment.

High dose oseltamivir, through its effect on lowering lung viral load, significantly reduced the levels of expression of TNF, IL-1β, IL-12p40 and CCL5 (Fig. 6 *M, N, P* and *R*). The combined treatment effectively dampened lung inflammation as early as 2 days after the initiation of treatment through reductions in the mRNA levels of TNF, IL-1β, IL-6, IL-12p40, CCL2, CCL5 and CXCL10 (Fig. 6 *M-S*).

Taken together, simultaneous targeting of virus and host inflammatory response by etanercept or STAT3 inhibitor is a viable strategy to treat severe IAV pneumonia, particularly when treatment needs to be initiated late after the onset of signs and symptoms. Significantly, combined treatment with oseltamivir and either of the anti-inflammatory drugs dampened an overlapping set of inflammatory factors that included TNF, IL-1β, IL-6, IL-12p40, CCL2, CCL5, and CXCL10.

## Discussion

Pneumonia is a severe complication caused by inflammation of the lungs due to infection with diverse viral pathogens that often results in respiratory failure and death. Seasonal and pandemic influenza viruses (1-4), variola virus (agent of smallpox) (35) and severe acute respiratory syndrome coronavirus 2 (SARS-CoV-2) (36) are leading examples. Pneumonia is one of the most common and life-threatening complications of IAV infection (2). Globally, nearly 1 billion people are infected with seasonal influenza annually, with 3-5 million cases of severe illness and 300,000-650,000 deaths (37, 38). In 2017 alone, 145,000 deaths and about 9.5 million hospitalizations were attributed to influenza-associated lower respiratory tract infection, including pneumonia (39).

There are no specific treatments for influenza pneumonia other than hospitalization and supportive care. Antivirals against IAV are ineffective against severe influenza and pneumonia if treatment is not initiated within 48 hours of disease symptoms. Most individuals do not seek medical attention within this timeframe (Choi et al, 2018). There is, therefore, an urgent need to advance therapies that specifically treat severe IAV post-onset of symptoms. This study reports an efficacious approach to treating influenza pneumonia by simultaneous treatment with an antiviral and an anti-inflammatory drug.

Strategies that target the virus alone with antivirals have shown limited clinical efficacy in treating IAV infection, especially when treatment is initiated late during the course of illness. Oseltamivir, the most commonly used anti-IAV drug, is only effective in reducing morbidity, hospital admissions or disease complications when treatment is commenced within 48 h of onset of symptoms (5). The ineffectiveness of antivirals in ameliorating pneumonia or reducing morbidity late after onset of disease symptoms is not unique to IAV infection. Several studies, including our own, have shown that cidofovir or tecovirimat (Tpoxx), antiviral agents that are effective in treating orthopoxvirus (OPXV) infections, are only partially protective if administered late after disease onset (33, 40-42). Similarly, antiviral drugs for the herpes simplex virus (acyclovir and valacyclovir) are effective only when administered within 72 h of lesion appearance (43). In patients hospitalized with SARS-CoV-2 infection, antiviral drugs, including remdesivir, lopinavir, and interferon were found to have little or no effect in reducing disease progression and mortality (44).

On the other hand, although strategies to limit or dampen inflammation during IAV infection have shown potential benefits, evidence on clinical benefits from the use of only anti-inflammatory drugs is inconclusive, particularly in hospitalized patients or when the treatment initiation is delayed (14). Corticosteroids, widely used as anti-inflammatory agents, were ineffective in preventing mortality of IAV H5N1 infected mice when treatment was initiated late after the onset of disease signs (45, 46). Other important regulators of inflammation, including peroxisome proliferator-activated receptor agonist and cyclooxygenase inhibitors, showed no survival benefits in H5N1 IAV infected mice when administered 48 h p.i. (47). In contrast, immunomodulatory agents targeting sphingosine-1-phosphate receptor agonist (48), C-C chemokine receptor 2 (49), or AMP-activated protein kinase (50) protected mice against lethal influenza virus challenge. These agents were administered as prophylactics or very early after virus inoculation. Such treatment regimens have a minimal translatory application, especially since patients often seek medical advice after the onset of symptoms or once they have developed pneumonia.

Both high viral load and high levels of inflammatory factors (cytokines, chemokines and transcription factors) increase the risk of illness severity and mortality after the onset of disease signs in H1N1 or H5N1 IAV infections in mice and humans (Fig. S1)(1, 9, 51, 52). That provided a rationale for the simultaneous targeting of virus replication and host inflammatory response as an approach to reduce morbidity and mortality in IAV infected mice. We co-administered oseltamivir and anti-inflammatory drugs (etanercept or SI-301) to reduce lung viral load and pathology and improve survival rates in mice with H1N1 influenza pneumonia.

We first focused on TNF, which is produced in the early phases of IAV infection and is associated with illness severity and morbidity in mice (53), swine (54) and human (55) influenza. Anti-TNF drugs are used widely to treat immune-mediated inflammatory diseases, including rheumatoid arthritis, Crohn’s disease, and psoriatic arthritis (56). However, in contrast to treating chronic inflammatory diseases, our results show that the effective treatment of IAV pneumonia would require multiple doses of etanercept. We have also made a similar finding in a mouse model of OPXV pneumonia (33), indicating that the levels of TNF produced in the lung during acute respiratory viral infection might be different from those produced during chronic inflammatory disease and as a result, requires multiple doses of etanercept.

Combined treatment with etanercept and oseltamivir, late after the onset of disease signs, reduced morbidity, viral load, and lung pathology in IAV-infected mice through downregulation of inflammatory factors. These included TNF, IL-1β, IL-6, IL-12p40, CCL2, CCL5 and CXCL10. It also reduced the activation of STAT3. Etanercept or oseltamivir monotherapy had limited clinical benefits given that they reduced the mRNA levels of some inflammatory factors but not to the extent of the combined treatment regimen. In a previous study, Shi and colleagues reported that etanercept treatment 2 h p.i. with H1N1 IAV protected mice from otherwise lethal infection (57). Our study evaluated the efficacy of combined therapy in a realistic timeframe that closely reflects timeframes when patients present at hospitals. Even before it has developed, treating lung inflammation in humans (even before it has developed) from the day of infection is neither feasible nor practical. However, if exposure to virulent or pandemic strains of IAV is identified very early, then anti-IAV drugs, at a standard recommended dose of 75 mg twice daily, would effectively minimize the risk of severe disease, morbidity and mortality (22). We found that a higher dose of oseltamivir at 150 mg/kg/day, but not a standard mouse dose of 40 mg/kg/day (58) effectively reduced the lung IAV load. A higher dose of oseltamivir should be considered for evaluation in a clinical setting, particularly in the milieu of severe lung inflammation when the treatment initiation might be delayed.

We made a similar finding when the STAT3 signaling pathway was targeted instead of the TNF/NF-κB pathway to reduce lung inflammation. Combined treatment with S3I-201 and oseltamivir reduced viral load, disease signs and lung pathology in IAV-infected mice through the downregulation of the same set of cytokines and chemokines as combined treatment with etanercept and oseltamivir, i.e., TNF, IL-1β, IL-6, IL-12p40, CCL2, CCL5 and CXCL10. These cytokines and chemokines enhance acute phase signaling, recruit inflammatory cells, including neutrophils, monocytes, and T lymphocytes to the site of infection, and trigger secondary cytokine production, resulting in lung inflammation and pathology (7). Thus, blockading cytokine(s) or cytokine signaling pathways combined with antiviral treatment will be expected to reduce leukocyte recruitment into the lung. Indeed, the combined treatment using either etanercept or SI-301 significantly reduced leukocyte migration to the lungs as evident histologically. In ECTV-infected mice, combined treatment with etanercept and cidofovir reduced recruitment of inflammatory monocytes to the lung (33). Inflammatory monocytes produce cytokines like IL-1, TNF, IL-6, CCL2 and CXCL10.

Various stimuli, including viral infection and inflammatory cytokines, activate the NF-κB and STAT3 signaling pathways. TNF and IL-1 activate NF-κB (59), which enhances cytokine expression, including IL-6, which is a potent inducer of STAT3 activation (13, 60, 61). NF-κB and STAT3 cooperatively regulate the expression of several inflammatory cytokines, such as IL-6, CCL2, CCL5, IL-8, and IL-17 (62, 63). A recent study evaluating global chromatin binding has revealed more than 36,000 *cis*-regulatory regions that can potentially bind to both STAT3 and NF-κB (64). NF-κB and STAT3 can collaboratively induce their target gene expression through direct physiological interaction or cooperative binding at a subset of gene promoters/enhancers (62, 64, 65).

Our results are in agreement with several other studies that have evaluated the therapeutic potential of adjunctive anti-inflammatory drug interventions in severe respiratory diseases. Corticosteroids in combination with antiviral agents effectively alleviate the 2009 pandemic H1N1 influenza-associated pneumonia (66). Similarly, Zheng *et al*. reported on the benefit of adjunctive therapy in an experimental mouse study where treatment was initiated late at 48 h after H5N1 IAV inoculation (47). They demonstrated amelioration of lung pathology and improved survival rates in virus-infected mice coadministration with cyclooxygenase inhibitors (mesalamine and celecoxib) and zanamivir. Besides IAV pneumonia, respiratory OPXV (26) and SARS-CoV-2 pneumonia (67, 68) are also associated with high viral load and dysregulated inflammation and may benefit from combined antiviral and anti-inflammatory treatment approaches. We have recently shown therapeutic efficacy of combined cidofovir (a viral DNA polymerase inhibitor) and etanercept or STAT3 inhibitor treatment in ameliorating lung pathology and protecting mice from lethal OPXV pneumonia (33). Similarly, Kalil et al. demonstrated superior effects of combination treatment with baricitinib, a Janus kinase inhibitor, and remdesivir, RNA-dependent RNA polymerase inhibitor, over remdesivir monotherapy in improving the clinical status of hospitalized COVID-19 patients (69).

Excessive early inflammatory cytokine/chemokine responses and leukocyte recruitment can predictict poor prognosis and poor clinical outcomes in IAV infections (1, 9, 10). Many inflammatory factors are responsible for leukocyte recruitment into the lungs (8, 11, 12). Our results indicated that targeting just one inflammatory cytokine (TNF) or a cytokine signaling pathway (STAT3) is sufficient to ameliorate lung pathology, reduce leukocyte infiltration, and confer protection from an otherwise lethal disease when treated combined with oseltamivir. An important finding in this study is that either etanercept or SI-301 treatment combined with oseltamivir dampened the same set of inflammatory cytokines/chemokines, i.e., TNF, IL-1β, IL-6, IL-12p40, CCL2, CCL5, and CXCL10. We believe that STAT3 inhibition will be more appropriate in individuals who cannot be treated with etanercept due to contraindications to the drug or when TNF may not be the driver of lung inflammation. Indeed, in a model of respiratory OPVX infection in TNF^-/-^ mice, combined treatment with cidofovir and SI-301 effectively reduced viral load and lung pathology (33).

In summary, combined treatment targeting virus and TNF/NF-κB or STAT3 pathways reduces viral load, clinical illness, and lung pathology in IAV infected mice through downregulation of inflammatory cytokines and chemokines, many of which are implicated in disease severity and lung pathology caused by other respiratory viruses, including OPXV (33) and coronaviruses (70). Therefore, the focus of clinical management of patients with severe viral pneumonia, associated with high viral load and dysregulated inflammation, should be on effective control of both viral replication and inflammatory cytokine and chemokine responses by the combined antiviral and anti-inflammatory drug treatment approach. More patient-oriented clinical research should be undertaken to test this treatment approach in severe respiratory diseases, in particular those caused by viruses of pandemic potential, which includes IAV and SARS-CoV-2. In the latter case, treatment efficacy would significantly increase when antivirals specific to the SARS-CoV-2 are available.

## Materials and Methods

### Animal Ethics Statement

Animal experiments were performed in accordance with protocols approved by the Animal Ethics Committee of the University of Tasmania (UTAS) (Protocol number A0016372) and the Animal Ethics and Experimentation Committee of the Australian National University (ANU) (Protocol numbers A2011/011 and A2014/018).

### Mice

C57BL/6J wild-type (WT) female mice, bred under specific pathogen-free conditions, were obtained from the Australian Phenomics Facility (APF), ANU, Canberra, the Cambridge Farm Facility, UTAS, Tasmania or the Animal Resource Centre, Western Australia, Australia. In addition to WT mice, TNF deficient (TNF^-/-^) (71) and triple mutant (TM or mTNF^Δ/Δ^.TNFRI^-/-^.II^-/-^, expressing only mTNF but lacking sTNF and TNFRs) (27) mice, bred at the APF, ANU were used in this study. One week prior to start of experiments, mice were transferred to the virus suite and allowed to acclimatize. Mice were used at 6-12 weeks of age.

### Cell Lines and Viruses

Madin-Darby Canine Kidney (MDCK) cells (ATTC No. CCL-34) were cultured in Dulbecco’s Modified Eagle Medium (DMEM) supplemented with 2mM L-glutamine (Sigma-Aldrich), antibiotics (1x PSN; 50 U/mL penicillin, 50 μg/mL streptomycin, and 100 μg/mL neomycin) (Sigma-Aldrich), and 10 % heat inactivated fetal bovine serum (FBS). This will be referred to as cell growth medium. Cell cultures were maintained at 37°C in a 5% CO_2_ atmosphere.

Stocks of Influenza A virus (IAV) H1N1 (A/PR/8/34) strain were propagated in 10-day old, specific-pathogen-free embryonated chicken eggs, and viral titers were determined in MDCK cells using plaque assay or median tissue culture infectious dose (TCID_50_) assay described elsewhere (72).

### Virus Infection, Animal Weights, and Clinical Scores

Age-matched mice were anesthetized with an isoflurane (UTAS) at 5% for induction and 2% for maintenance intranasally (i.n.) using the Stinger Streamline Rodent/Exotics Anesthetic Gas Machine (Advanced Anesthesia Specialists) or with tribromoethanol (ANU) at 160-240 mg/kg through the intraperitoneal (i.p.) route. Mice were infected with 3000 plaque forming unit (PFU) of IAV, housed in individually ventilated cages under biological safety level 2 containment facilities and monitored daily; weighed and scored for clinical signs of illness (scores ranged from 0 to 3 for each of the five clinical parameters, condition of hair coat, posture, breathing, lacrimation and nasal discharge, and activity and behavior) as described elsewhere (26, 27). For ethical reasons and as required by the animal ethics protocols, mice that were severely moribund with a clinical score of ≥ 10 and/or a body weight loss of ≥ 20% (UTAS) or 25% (ANU) were killed by CO_2_ asphyxiation, lung tissues collected for subsequent analyses and animals considered dead the following day.

### Drug Treatments

Different groups of mice were administered with 100 µl of oseltamivir (Tamiflu; Roche) diluted in phosphate buffered saline (PBS) via oral gavage (o.g.) at 150 mg/kg (once daily) or 20 mg/kg (twice daily), etanercept (Enbrel; Pfizer Inc.) i.p. at 2.5 mg/kg diluted in PBS or S3I-201 (STAT3 inhibitor VI, Sigma-Aldrich, cat. no. 573102) (i.p.) at 5 mg/kg after the onset of disease signs. S3I-201 was first dissolved in DMSO at 200 mg/mL and then further diluted in PBS to a working stock solution of 1 mg/mL.

### Plaque Assay for Virus Quantification

Viral titers were determined as virus plaque forming units (PFU) per gram of lung tissue samples using a plaque assay as described elsewhere (72). Briefly, homogenized lung samples were serially diluted (10-fold) and inoculated into the confluent MDCK monolayer in a 6-well tissue culture plate. After 1 h incubation, inoculum was removed and the monolayer was covered with the agar overlay. After 4 days of incubation, the agar overlay was removed and the cells were fixed with 10% formalin followed by staining with 0.1% crystal violet. Viral titers were calculated as PFU/g. For details, see Supplementary Information, Materials and Methods.

### TCID_50_ Assay for Virus Quantification

TCID_50_ assay for IAV quantification was previously described elsewhere (72). Briefly, serial dilutions of lung homogenates were inoculated into the MDCK cell monolayer in a 96-well tissue culture plate. After 1 h of virus adsorption, inoculum was removed and the infected monolayer was incubated in the virus growth medium at 37 °C, 5% CO_2_ for 4 days. The cells were then fixed with 10% formalin and stained with 0.1% crystal violet to visualize virus induced cell cytopathic effect (CPE). We calculated the viral titer as TCID_50_/g using the Reed-Muench method (73). For details, see Supplementary Information, Materials and Methods.

### Lung Histopathological Examination

Lung tissue was sectioned and stained with Hematoxylin and eosin (H&E) and assessed for histopathology using a semi-quantitative scoring system as described elsewhere (26, 27). For more details, see Supplementary Information, Materials and Methods.

### RNA Extraction, cDNA Generation and quantitative reverse transcription polymerase chain reaction (qRT-PCR)

RNA was extracted from the lung tissue homogenized in TRIzol solution (ThermoFisher Scientific, cat. no. 15596026) as described elsewhere (26, 27), and cDNA was synthesized using RevertAid first strand cDNA synthesis kit (ThermoFisher Scientific, cat. no. K1622). PowerUp SYBR Green Master Mix (ThermoFisher Scientific, cat. no. A25742) was used to measure mRNA transcripts of cytokines/chemokines levels using quantitative real-time PCR (qRT-PCR). Recorded cycle threshold values were normalized to the housekeeping gene Ubiquitin C (UBC). Details on the procedure and primers used are presented in Supplementary Information, Materials and Methods.

### Protein Extraction and Western Blot Analysis

Total protein extraction from the lung tissue and western blot analysis was undertaken as described elsewhere (26). Details on the procedure and antibodies used are included in Supplementary Information, Materials and Method*s*.

### Statistical Analysis

Statistical analyses of experimental data, as indicated in the legend to each figure, were performed using appropriate tests to compare results using GraphPad Prism 9 (GraphPad Software, Inc.). A value of P < 0.05 was taken to be significant: *, p < 0.05; **, p < 0.01; ***, p <0.001 and ****, p <0.0001.

## Acknowledgments

This work was supported by grants from the National Health and Medical Research Council of Australia (Grant 1007980) and the Medical Protection Society of Tasmania Inc., Tasmania, Australia. The funders had no role in study design, data collection and interpretation, or the decision to submit the work for publication. We gratefully acknowledge Professor Jane Dahlstrom, Canberra Hospital for helping us develop the lung histopathology scoring system. We are grateful to the UTAS Animal Services Division and the ANU Phenomics Facility for the care and breeding of mice. P.P. was the recipient of a Tasmanian School of Medicine Research Scholarship through the University of Tasmania.

## References

1. M. D. de Jong et al., Fatal outcome of human influenza A (H5N1) is associated with high viral load and hypercytokinemia. Nat Med 12, 1203–1207 (2006).

2. J. Rello, A. Pop-Vicas, Clinical review: primary influenza viral pneumonia. Critical care (London, England) 13, 235 (2009).

3. J. Rello et al., Intensive care adult patients with severe respiratory failure caused by Influenza A (H1N1)v in Spain. Critical care (London, England) 13, R148 (2009).

4. M. B. Rothberg, S. D. Haessler, R. B. Brown, Complications of viral influenza. The American journal of medicine 121, 258–264 (2008).

5. T. Jefferson et al., Neuraminidase inhibitors for preventing and treating influenza in healthy adults and children. The Cochrane database of systematic reviews 2014, Cd008965 (2014).

6. S. H. Choi et al., Late diagnosis of influenza in adult patients during a seasonal outbreak. Korean J Intern Med 33, 391–396 (2018).

7. J. R. Tisoncik et al., Into the eye of the cytokine storm. Microbiology and Molecular Biology Reviews 76, 16–32 (2012).

8. N. L. La Gruta, K. Kedzierska, J. Stambas, P. C. Doherty, A question of self-preservation: immunopathology in influenza virus infection. Immunology and cell biology 85, 85–92 (2007).

9. D. Kobasa et al., Aberrant innate immune response in lethal infection of macaques with the 1918 influenza virus. Nature 445, 319–323 (2007).

10. C. M. Oshansky et al., Mucosal Immune Responses Predict Clinical Outcomes during Influenza Infection Independently of Age and Viral Load. American Journal of Respiratory and Critical Care Medicine 189, 449–462 (2014).

11. T. J. Schall, K. B. Bacon, Chemokines, leukocyte trafficking, and inflammation. Current opinion in immunology 6, 865–873 (1994).

12. R. Alon et al., Leukocyte trafficking to the lungs and beyond: lessons from influenza for COVID-19. Nature reviews. Immunology 21, 49–64 (2021).

13. P. Pandey, G. Karupiah, Targeting tumour necrosis factor to ameliorate viral pneumonia. The FEBS journal (2021).

14. Q. Liu, Y. H. Zhou, Z. Q. Yang, The cytokine storm of severe influenza and development of immunomodulatory therapy. Cellular & molecular immunology 13, 3–10 (2016).

15. R. L. Peper, H. Van Campen, Tumor necrosis factor as a mediator of inflammation in influenza A viral pneumonia. Microbial pathogenesis 19, 175–183 (1995).

16. S. E. Belisle et al., Genomic profiling of tumor necrosis factor alpha (TNF-alpha) receptor and interleukin-1 receptor knockout mice reveals a link between TNF-alpha signaling and increased severity of 1918 pandemic influenza virus infection. Journal of virology 84, 12576–12588 (2010).

17. F. G. Hayden et al., Local and systemic cytokine responses during experimental human influenza A virus infection. Relation to symptom formation and host defense. The Journal of clinical investigation 101, 643–649 (1998).

18. B. R. B. Pires, R. Silva, G. M. Ferreira, E. Abdelhay, NF-kappaB: Two Sides of the Same Coin. Genes 9 (2018).

19. M. S. Hayden, S. Ghosh, Regulation of NF-κB by TNF family cytokines. Semin Immunol 26, 253–266 (2014).

20. A. D. Watts et al., A casein kinase I motif present in the cytoplasmic domain of members of the tumour necrosis factor ligand family is implicated in ‘reverse signalling’. The EMBO journal 18, 2119–2126 (1999).

21. G. Eissner, W. Kolch, P. Scheurich, Ligands working as receptors: reverse signaling by members of the TNF superfamily enhance the plasticity of the immune system. Cytokine & growth factor reviews 15, 353–366 (2004).

22. DTB, Oseltamivir for influenza. Drug and therapeutics bulletin 40, 89–91 (2002).

23. CDC (2021) Influenza Antiviral Medications: Summary for Clinicians. (Centers for DIsease and Prevention).

24. E. De Clercq, Antiviral agents active against influenza A viruses. Nature reviews. Drug discovery 5, 1015–1025 (2006).

25. S. Ramiro et al., Combination therapy for pain management in inflammatory arthritis (rheumatoid arthritis, ankylosing spondylitis, psoriatic arthritis, other spondyloarthritis). The Cochrane database of systematic reviews 10.1002/14651858.CD008886.pub2, Cd008886 (2011).

26. M. J. Tuazon Kels et al., TNF deficiency dysregulates inflammatory cytokine production, leading to lung pathology and death during respiratory poxvirus infection. Proceedings of the National Academy of Sciences of the United States of America 117, 15935–15946 (2020).

27. Z. Al Rumaih et al., Poxvirus-encoded TNF receptor homolog dampens inflammation and protects from uncontrolled lung pathology during respiratory infection. Proceedings of the National Academy of Sciences 117, 26885–26894 (2020).

28. Z. Kaymakcalan et al., Comparisons of affinities, avidities, and complement activation of adalimumab, infliximab, and etanercept in binding to soluble and membrane tumor necrosis factor. Clinical immunology (Orlando, Fla.) 131, 308–316 (2009).

29. U. Meusch, M. Rossol, C. Baerwald, S. Hauschildt, U. Wagner, Outside-to-inside signaling through transmembrane tumor necrosis factor reverses pathologic interleukin-1beta production and deficient apoptosis of rheumatoid arthritis monocytes. Arthritis and rheumatism 60, 2612–2621 (2009).

30. D. Damjanovic et al., Negative regulation of lung inflammation and immunopathology by TNF-α during acute influenza infection. The American journal of pathology 179, 2963–2976 (2011).

31. R. Dutkowski et al., Safety and pharmacology of oseltamivir in clinical use. Drug safety 26, 787–801 (2003).

32. P. Ward, I. Small, J. Smith, P. Suter, R. Dutkowski, Oseltamivir (Tamiflu) and its potential for use in the event of an influenza pandemic. The Journal of antimicrobial chemotherapy 55 Suppl 1, i5–i21 (2005).

33. P. Pandey et al., Targeting ectromelia virus and TNF/NF-κB or STAT3 signaling for effective treatment of viral pneumonia. Proceedings of the National Academy of Sciences of the United States of America In Press (2022).

34. G. Karupiah, V. Panchanathan, I. G. Sakala, G. Chaudhri, Genetic resistance to smallpox: lessons from mousepox. Novartis Foundation symposium 281, 129–136; discussion 136-140, 208-129 (2007).

35. P. J. Ross, A. Seaton, H. M. Foreman, W. H. Morris Evans, Pulmonary calcification following smallpox handler’s lung. Thorax 29, 659–665 (1974).

36. P. Zhou et al., A pneumonia outbreak associated with a new coronavirus of probable bat origin. Nature 579, 270–273 (2020).

37. A. D. Iuliano et al., Estimates of global seasonal influenza-associated respiratory mortality: a modelling study. Lancet (London, England) 391, 1285–1300 (2018).

38. O. World Health, Global influenza strategy 2019-2030 (World Health Organization, Geneva, 2019).

39. G. I. Collaborators, Mortality, morbidity, and hospitalisations due to influenza lower respiratory tract infections, 2017: an analysis for the Global Burden of Disease Study 2017. The Lancet. Respiratory medicine 7, 69–89 (2019).

40. D. C. Quenelle, D. J. Collins, E. R. Kern, Efficacy of multiple-or single-dose cidofovir against vaccinia and cowpox virus infections in mice. Antimicrob Agents Chemother 47, 3275–3280 (2003).

41. M. Bray et al., Cidofovir protects mice against lethal aerosol or intranasal cowpox virus challenge. The Journal of infectious diseases 181, 10–19 (2000).

42. A. T. Russo et al., Effects of Treatment Delay on Efficacy of Tecovirimat Following Lethal Aerosol Monkeypox Virus Challenge in Cynomolgus Macaques. The Journal of infectious diseases 218, 1490–1499 (2018).

43. C. Cernik, K. Gallina, R. T. Brodell, The Treatment of Herpes Simplex Infections: An Evidence-Based Review. Archives of Internal Medicine 168, 1137–1144 (2008).

44. W. S. T. Consortium, Repurposed Antiviral Drugs for Covid-19 — Interim WHO Solidarity Trial Results. New England Journal of Medicine 384, 497–511 (2020).

45. T. Xu et al., Effect of dexamethasone on acute respiratory distress syndrome induced by the H5N1 virus in mice. The European respiratory journal 33, 852–860 (2009).

46. R. Salomon, E. Hoffmann, R. G. Webster, Inhibition of the cytokine response does not protect against lethal H5N1 influenza infection. Proceedings of the National Academy of Sciences of the United States of America 104, 12479–12481 (2007).

47. B.-J. Zheng et al., Delayed antiviral plus immunomodulator treatment still reduces mortality in mice infected by high inoculum of influenza A/H5N1 virus. Proceedings of the National Academy of Sciences 105, 8091–8096 (2008).

48. K. B. Walsh et al., Suppression of cytokine storm with a sphingosine analog provides protection against pathogenic influenza virus. Proceedings of the National Academy of Sciences of the United States of America 108, 12018–12023 (2011).

49. K. L. Lin, S. Sweeney, B. D. Kang, E. Ramsburg, M. D. Gunn, CCR2-antagonist prophylaxis reduces pulmonary immune pathology and markedly improves survival during influenza infection. Journal of immunology (Baltimore, Md.: 1950) 186, 508–515 (2011).

50. C. E. Moseley, R. G. Webster, J. R. Aldridge, Peroxisome proliferator-activated receptor and AMP-activated protein kinase agonists protect against lethal influenza virus challenge in mice. Influenza and other respiratory viruses 4, 307–311 (2010).

51. A. C. Boon et al., H5N1 influenza virus pathogenesis in genetically diverse mice is mediated at the level of viral load. mBio 2 (2011).

52. K. K. To et al., Delayed clearance of viral load and marked cytokine activation in severe cases of pandemic H1N1 2009 influenza virus infection. Clinical infectious diseases: an official publication of the Infectious Diseases Society of America 50, 850–859 (2010).

53. T. Hussell, A. Pennycook, P. J. Openshaw, Inhibition of tumor necrosis factor reduces the severity of virus-specific lung immunopathology. European journal of immunology 31, 2566–2573 (2001).

54. K. Van Reeth, S. Van Gucht, M. Pensaert, Correlations between lung proinflammatory cytokine levels, virus replication, and disease after swine influenza virus challenge of vaccination-immune pigs. Viral Immunol 15, 583–594 (2002).

55. R. S. Fritz et al., Nasal cytokine and chemokine responses in experimental influenza A virus infection: results of a placebo-controlled trial of intravenous zanamivir treatment. The Journal of infectious diseases 180, 586–593 (1999).

56. L. C. Silva, L. C. Ortigosa, G. Benard, Anti-TNF-α agents in the treatment of immune-mediated inflammatory diseases: mechanisms of action and pitfalls. Immunotherapy 2, 817–833 (2010).

57. X. Shi et al., Inhibition of the inflammatory cytokine tumor necrosis factor-alpha with etanercept provides protection against lethal H1N1 influenza infection in mice. Critical care (London, England) 17, R301 (2013).

58. D. F. Smee, M. H. Wong, K. W. Bailey, R. W. Sidwell, Activities of oseltamivir and ribavirin used alone and in combination against infections in mice with recent isolates of influenza A (H1N1) and B viruses. Antiviral chemistry & chemotherapy 17, 185–192 (2006).

59. T. Lawrence, The nuclear factor NF-kappaB pathway in inflammation. Cold Spring Harbor perspectives in biology 1, a001651 (2009).

60. J. G. Bode, U. Albrecht, D. Häussinger, P. C. Heinrich, F. Schaper, Hepatic acute phase proteins--regulation by IL-6-and IL-1-type cytokines involving STAT3 and its crosstalk with NF-κB-dependent signaling. European journal of cell biology 91, 496–505 (2012).

61. A. Oeckinghaus, M. S. Hayden, S. Ghosh, Crosstalk in NF-κB signaling pathways. Nat Immunol 12, 695–708 (2011).

62. J. Yang et al., Unphosphorylated STAT3 accumulates in response to IL-6 and activates transcription by binding to NFkappaB. Genes & development 21, 1396–1408 (2007).

63. S. I. Grivennikov, M. Karin, Dangerous liaisons: STAT3 and NF-kappaB collaboration and crosstalk in cancer. Cytokine Growth Factor Rev 21, 11–19 (2010).

64. Z. Ji, L. He, A. Regev, K. Struhl, Inflammatory regulatory network mediated by the joint action of NF-kB, STAT3, and AP-1 factors is involved in many human cancers. Proceedings of the National Academy of Sciences 116, 9453–9462 (2019).

65. H. Lee et al., Persistently activated Stat3 maintains constitutive NF-kappaB activity in tumors. Cancer Cell 15, 283–293 (2009).

66. K. Kudo et al., Systemic corticosteroids and early administration of antiviral agents for pneumonia with acute wheezing due to influenza A(H1N1)pdm09 in Japan. PloS one 7, e32280 (2012).

67. J. Fajnzylber et al., SARS-CoV-2 viral load is associated with increased disease severity and mortality. Nature communications 11, 5493 (2020).

68. C. Huang et al., Clinical features of patients infected with 2019 novel coronavirus in Wuhan, China. The Lancet 395, 497–506 (2020).

69. A. C. Kalil et al., Baricitinib plus Remdesivir for Hospitalized Adults with Covid-19. New England Journal of Medicine 384, 795–807 (2020).

70. V. A. Ryabkova, L. P. Churilov, Y. Shoenfeld, Influenza infection, SARS, MERS and COVID-19: Cytokine storm – The common denominator and the lessons to be learned. Clinical Immunology 223, 108652 (2021).

71. H. Körner et al., Distinct roles for lymphotoxin-α and tumor necrosis factor in organogenesis and spatial organization of lymphoid tissue. European journal of immunology 27, 2600–2609 (1997).

72. A. L. Balish, J. M. Katz, A. I. Klimov, Influenza: propagation, quantification, and storage. Current protocols in microbiology 29, 15G. 11.11-15G. 11.24 (2013).

73. L. J. Reed, H. Muench, A simple method of estimating fifty percent endpoints. American Journal of Epidemiology 27, 493–497 (1938).

