## Supplementary data and information for "Simultaneous therapeutic targeting of inflammation and virus ameliorates influenza pneumonia and protects from morbidity and mortality"

**This file includes:**
Supplementary text
Supplementary Figures S1 to S6 and legends
References

### Supplementary Information Text

### Methods

#### Plaque assay for virus quantification

Plaque assay for IAV quantification was previously described elsewhere (1). Briefly, Madin-Darby Canine Kidney (MDCK) cells were seeded in a 6-well tissue culture plate at 1.2 × $10^6$ cells/well and incubated overnight in the cell growth media. On the day of infection, 2x Leibovitz L15 medium (ThermoFisher scientific, cat. no. 41300039) and 1.8 % low melting point agarose (Lonza, cat. no. 51101) were pre-warmed in a water bath at 37°C and 46°C, respectively until required. Each lung sample was then weighed and homogenized in serum free DMEM media using TissueLyser II (Qiagen, cat no. 85300). The homogenized tissue samples were then sonicated at 100W power in three 15-second bursts in a cup sonicator (Branson Sonic Power Company, Danbury, CT, USA) to further break up clumps of tissue and release virus. MDCK cell monolayers were washed with serum free DMEM medium leaving around 0.2 mL of media in each well. Six 10-fold serial dilutions of the homogenized lung samples were prepared in serum free DMEM supplemented with 1x PSN antibiotics and 1.5 µg/mL L-1-Tosylamide-2-phenylethyl chloromethyl ketone (TPCK)-trypsin (Sigma-Aldrich, cat. no. T1426). This will be referred to as virus growth medium. From each dilution, a 100 µl volume was added to the cell monolayer, which was then incubated for 1 h at 37°C with intermittent shaking every 15 mins. Pre-warmed 2x Leibovitz L15 medium (Invitrogen, cat. no. 41300039) and 1.8% low melting point agarose (Lonza, cat. no. 50101) were mixed in equal (1:1) ratio to prepare the agar overlay which was supplemented with TPCK- trypsin at 1.5 µg/mL. The infected monolayer was then covered with 3 mL agar overlay. After agar solidification, the tissue culture plates were incubated at 37°C, 5 % $CO_2$ for 4 days. The cells were next fixed with 10% formalin for 30 min at room temperature and the agar plug was removed using a sterile spatula. Finally, plaques were visualized by staining with 200 µl 0.1% crystal violet followed by rinsing with water. To determine the viral titer of the sample, plaques were counted and the following equation was used.

$$\text{PFU/g lung} = \frac{\text{No. of plaques X dilution factor X vol. of homogenization solvent (µL)}}{\text{vol. of viral suspension applied (µL) X weight of lung tissue (g)}}$$

**$TCID_{50}$ Assay for Virus Quantification**

$TCID_{50}$ assay for IAV quantification has been described elsewhere (1). Briefly, MDCK cells were seeded in a 96-well tissue culture plates at $2.5 \times 10^4$ cells per well and grown overnight in the cell growth media. Lungs were homogenized as described for the plaque assay. Eight 10-fold serial dilutions of lung homogenates were prepared in the virus growth medium and 25 μL of each dilution was inoculated onto cell monolayers in 10 replicates. After 1 h of virus adsorption, the inoculum was removed and the infected monolayer was incubated in virus growth medium at 37°C, 5% $CO_2$ for 4 days. The cells were then fixed with 10% formalin and stained with 0.1% crystal violet to visualize virus induced cell cytopathic effect (CPE). The viral titer was calculated as $TCID_{50}$/g lung tissue using the Reed-Muench method (2) and estimated the corresponding PFU using the conversion PFU = 0.7 $TCID_{50}$ (3).

**Histology and microscopic examination of lung pathology**

The left lung was dissected out and fixed in 10% neutral-buffered formalin at room temperature for 24 h, processed in the Leica ASP300S tissue processor, embedded in paraffin, 6 μm thick sagittal sections were cut and stained with H&E for analysis with a bright field microscope. A semi-quantitative scoring system developed previously in our laboratory as described elsewhere (4, 5) was used for assessing lung pathology. Blinded visual scoring of individual slides were done on a scale from 0 to 4 for each of the six criteria: parenchymal edema, perivascular edema, degree of bronchial epithelial necrosis, parenchymal inflammatory infiltrates, perivascular inflammatory infiltrates, and alveolar septal wall damage.

**RNA extraction and cDNA generation**

Lung tissue from individual mice was homogenized in 1 mL TRIzol reagent (ThermoFisher Scientific, cat. no. 15596026) using TissueLyser II (Qiagen, cat. no. 853000) as described elsewhere (4, 5). Genomic DNA traces were removed by treatment with RQ1 RNase-free DNase (Promega, cat. no. M6101) and the RNA concentration was determined using a Nanodrop ND-1000 spectrophotometer (ThermoFisher Scientific). cDNA was synthesized from 2 μg of RNA in a 20 μL volume using RevertAid first strand cDNA synthesis kit (ThermoFisher Scientific, cat. no. K1622) and incubated for 5 min at

25°C followed by 60 min at 42°C. cDNA samples were diluted with 20 µL nuclease-free water and were stored at -20°C until further use.

**Quantitative reverse transcription real-time polymerase chain reaction (qRT-PCR)**

qRT-PCR was performed using a 10 µL PCR reaction mixture prepared by mixing 5 µL 2X PowerUp SYBR green master mix (ThermoFisher Scientific, cat. no. A25742), 0.5 µM gene-specific forward and reverse primers (listed below in the table), 1 µL of cDNA template, and 3 µL nuclease-free water in the QuantStudio 3 (ThermoFisher Scientific). The following conditions were used: uracil-DNA glycosylase (UDG) activation at 50°C for 2 min, initial denaturation at 95°C for 2 min, and 40 cycles of denaturation and annealing/extension at 95°C for 3 sec and 60°C for 30 sec, respectively. The relative gene expression was calculated with the delta-delta Ct method (6) using ubiquitin C (UBC) for normalization. Results are reported as the fold-change relative to gene expression in mock-treated mice using the delta-delta Ct method (6).

List of primers used in this study are as follows:

| Gene | Primer sequence (5' - 3') | |
|---|---|---|
| **UBC** | Forward | AGGTCAAACAGGAAGACAGACGTA |
| | Reverse | TCACACCCAAGAACAAGCACA |
| **CCL2** | Forward | TTAAAAACCTGGATCGGAACCAA |
| | Reverse | GCATTAGCTTCAGATTTACGGGT |
| **CCL5** | Forward | GCTGCTTTGCCTACCTCTCC |
| | Reverse | TCGAGTGACAAACACGACTGC |
| **CXCL10** | Forward | TGAGTGGGACTCAAGGGATCC |
| | Reverse | TTCAAGCTTCCCTATGGCCC |
| **TNF** | Forward | ACTTCGGGGTGATCGGTCCCC |
| | Reverse | CCACTTGGTGGTTTGCTACGACGT |
| **IL-6** | Forward | TAGTCCTTCCTACCCCAATTTCC |
| | Reverse | TTGGTCCTTAGCCACTCCTTC |
| **IL-1β** | Forward | GCAACTGTTCCTGAACTCAACT |
| | Reverse | ATCTTTTGG GGT CCG TCA ACT |
| **IL-12p40** | Forward | AGACCCTGCCCATTGAACTG |
| | Reverse | CGGGTCTGGTTTGATGATGTC |

**Protein extraction and western blot analysis**

Whole cell lysate from lung tissue was prepared in 200 µL of RIPA buffer supplemented with 1X protease/phosphatase inhibitor cocktail (Cell Signalling Technology, cat. no. 5872S) using TissueLyser II as described elsewhere (4). Total protein was quantified using the fluorescence based EZQ protein quantification kit (ThermoFisher Scientific, cat. no. R33200). The stained protein spots were analyzed in a microplate reader (Spark multimode plate reader, Tecan Life Sciences) using excitation/emission settings of 485/590 nm. Protein concentrations were then calculated using the standard curve and normalized to 5 µg/µL.

For protein denaturation, a 25 µg protein lysate with an equal volume of 2X Laemmli buffer was incubated at 95$^{o}$C for 5 min. Denatured protein lysates were then separated by sodium dodecyl sulfate -polyacrylamide gel electrophoresis (SDS-PAGE) on Novex Tris-Glycine gels (ThermoFisher Scientific, cat. no XP00102) and transferred onto polyvinylidene fluoride (PVDF) membranes (ThermoFisher Scientific, cat. no IB24001) using iBlot 2 Gel Transfer Device (ThermoFisher Scientific) at the P0 preset template: 20 V for 1 min, 23 V for 4 min and 25 V for 2 min. Membrane was then blocked with 5% non-fat dry milk in Tris-buffered saline with Tween solution (TBST, 50 mM Tris-HCl, PH 8.0, 150 mM NaCl, and 0.1% Tween-20) for 1 h followed by incubation with primary antibody (listed in the table below) overnight at $4^{o}C$. The immunoblot was then incubated with a horseradish peroxidase (HRP)-coupled secondary antibody (listed in the table below) for 1 h at room temperature. Finally, the blot was developed using SuperSignal West Pico PLUS Chemiluminescent Substrate (ThermoFisher Scientific, cat. no. 34580) and visualized in a chemiluminescent imager (Amersham 600, Cytiva Life Sciences). Protein bands in the immunoblot were quantified with the ImageJ software (https://imagej.net/software/fiji/) as described elsewhere (7). Each target protein band was first normalized to β-actin band and expressed as a ratio of normalized protein band intensities relative to the mock treated control.

Primary and secondary antibodies used for western blotting

| Antibody | Isotype | Clone | Supplier | Catalog no. |
|---|---|---|---|---|
| I. Primary antibody | | | | |
| TNF | Armenian Hamster IgG | TN3.19-12 | Santa Cruz Biotechnology | sc-12744 |
| Phospho-STAT3 (Tyr705) | Rabbit IgG | D3A7 | Cell Signalling Technology | 9145 |
| Phospho-NF-κB p65 (Ser 536) | Rabbit IgG | T.849.2 | ThermoFisher Scientific | MA5-15160 |
| β-actin | Rabbit IgG | 13E5 | Cell Signalling Technology | 4970 |
| II. Secondary antibody | | | | |
| anti-rabbit IgG | Goat IgG | Polyclonal | Santa Cruz Biotechnology | sc-2004 |
| anti-Armenian hamster IgG | Goat IgG | Polyclonal | Santa Cruz Biotechnology | sc-2443 |

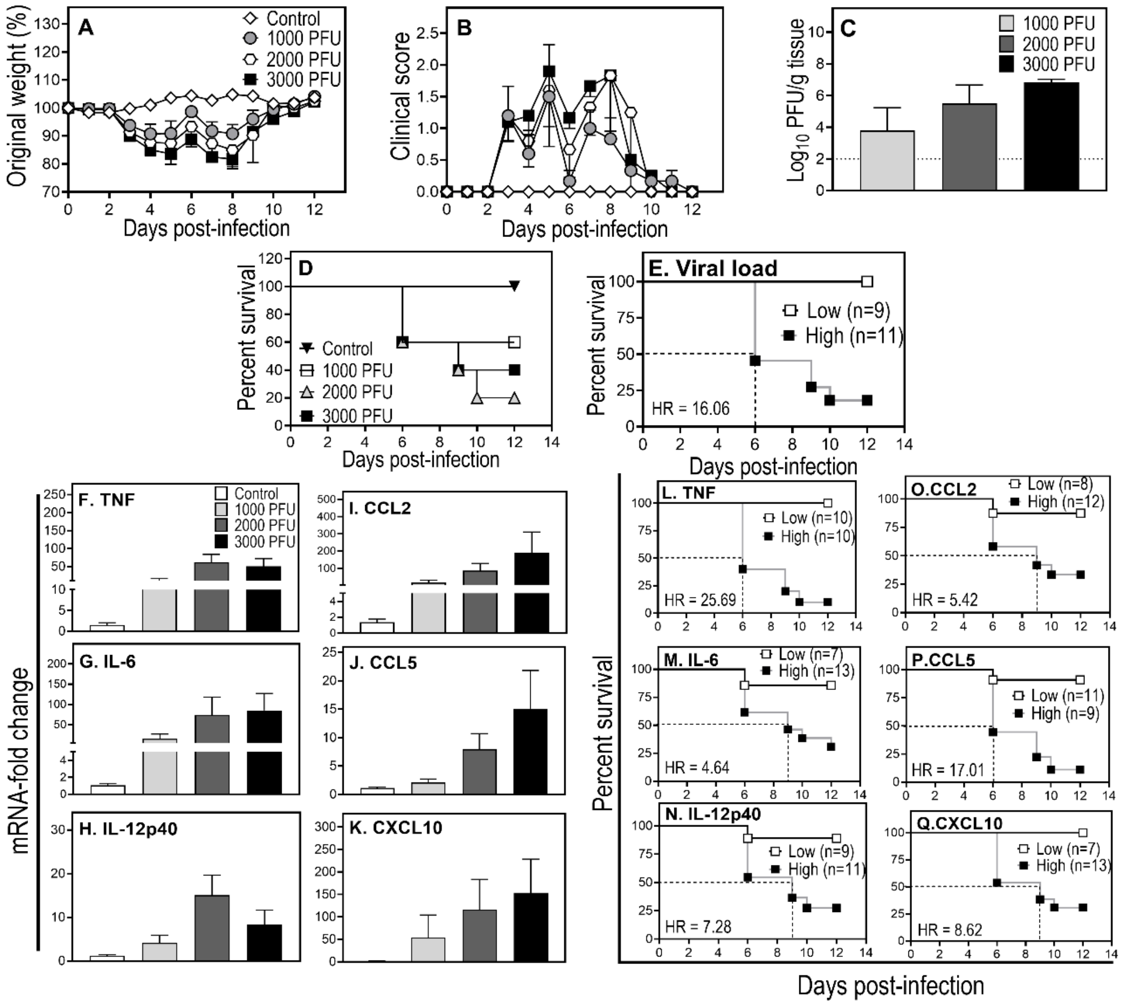


**Fig. S1. High lung viral load and cytokine/chemokine levels are associated with increased mortality of IAV-infected mice.** Age-matched groups of WT mice (n = 5) were infected with 1000, 2000, or 3000 PFU IAV i.n. Weight loss (A) and clinical scores (B) were assessed until day 12 p.i., when all the animals were killed and lungs collected for various analyses. Lung viral load (C) data were log-transformed and the detection limit of $TCID_{50}$ assay has been represented by a broken line. Survival data (D) were analyzed using log-rank (Mantel-Cox) test. The median survival times for mice infected with 1000, 2000 and 3000 PFU IAV were >12, 9 and 9 days, respectively compared with a median survival of >12 days for uninfected mice (log-rank test, uninfected vs 1000 PFU, $p = 0.134$; uninfected vs 2000 PFU, $p = 0.013$; uninfected vs 3000 PFU, $p = 0.049$). (E) Data from all groups were combined to evaluate survival rate in mice with low or high lung viral load (cut off threshold was $10^4$ PFU). (F-K) Gene transcript levels of cytokines and chemokines were evaluated using qPCR. Data are expressed as means ± SEM. (L-Q) Data from all groups combined to assess survival rate between mice having low and high cytokine/chemokine levels. The cutoff values that define high and low levels of cytokines and chemokines were chosen based on the highest level of that cytokine or chemokine among uninfected mice. Dotted lines indicate the median survival time (days). Survival distribution of mice between high and low lung viral titers (E) and cytokine expression levels (L-Q) were assessed by hazard ratio (HR). Numbers in parentheses indicate mouse numbers in high/low virus titer groups (panel E) and mouse numbers in high/low cytokine/chemokine levels (panels L-Q). In our hands, the 3000 PFU dose (used throughout the study) has been consistently lethal for C57BL/6 mice but was not the case in this experiment. Data shown are from a single experiment.

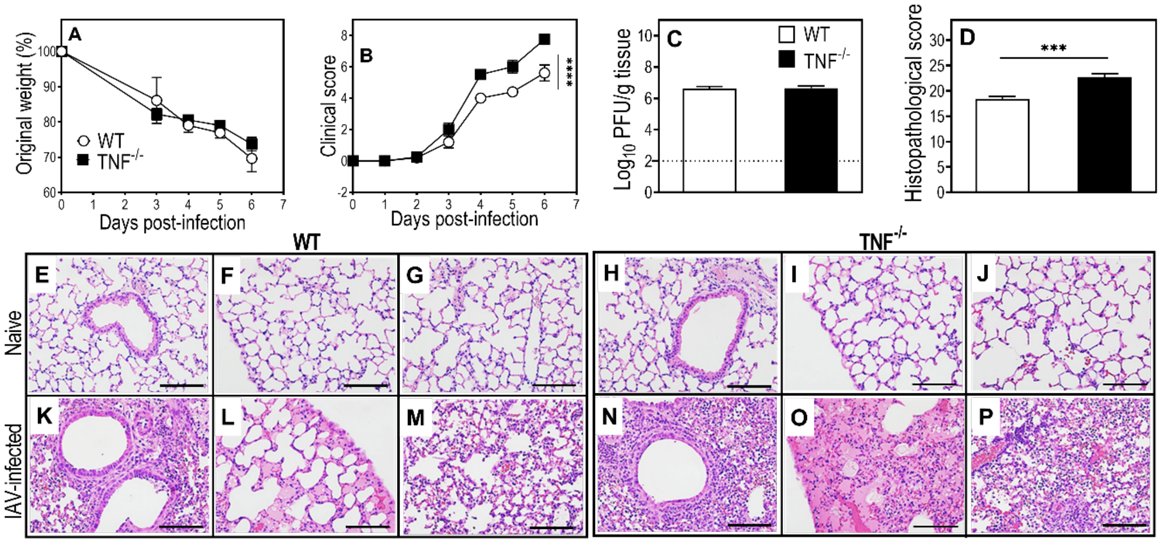


**Fig. S2. TNF deficiency exacerbates clinical signs and lung pathology in IAV-infected mice.** Age-matched groups of female WT and TNF$^{-/-}$ mice (n = 4 or 5) were infected with 3000 PFU IAV i.n. Weight loss and clinical scores (A and B) were assessed until day 6 p.i. when animals were killed. Lungs were collected for various analyses. Lung viral load (C) data were log-transformed and histopathological scores (D) were based on microscopic examination of lung histology H&E sections (E-P), which show that edema, leukocyte infiltration and alveolar septa damage are higher in the lungs of IAV-infected TNF$^{-/-}$ deficient mice compared to WT mice. H&E sections were examined using bright field microscope on all fields at 400x magnification. Data are expressed as means ± SEM and were analyzed using two-way ANOVA with Sidak's post-tests for (A) and (B) and unpaired t-test for (C) and (D). ***, $p \le 0.001$ and ****, $p \le 0.0001$. Broken line in panel C corresponds to the limit of virus detection. Bars in panels E-P correspond to 100 µm. Data shown are from a single experiment.

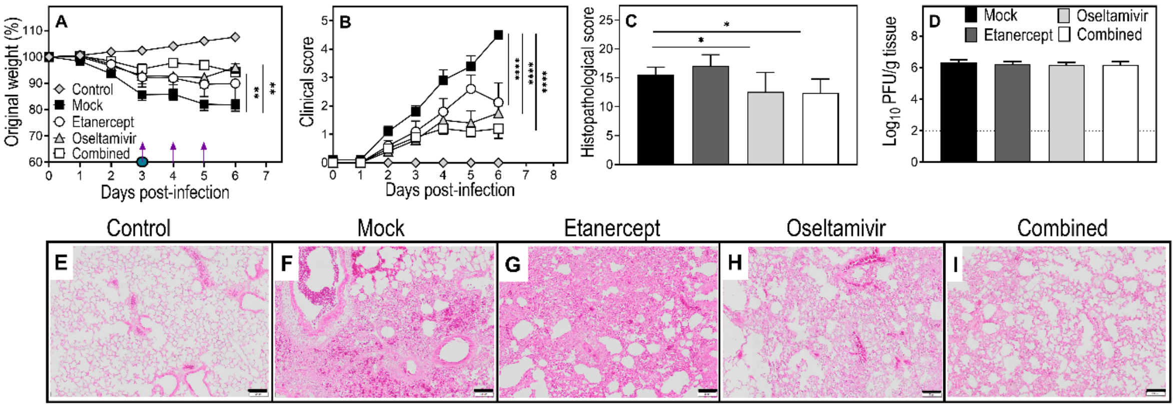


**Fig. S3. One dose of etanercept combined with standard dose oseltamivir reduces weight loss, clinical scores and lung pathology but not viral load in IAV-infected WT mice**. Age-matched groups of female WT mice (n = 5) were infected with 3000 PFU of IAV i.n. Animals were treated with etanercept, oseltamivir (20mg/kg, twice a day), or both drugs (combined) on day 3 p.i. Oseltamivir treatment was continued on days 4 and 5 p.i. as indicated in panel A, where a purple arrow and a filled blue circle symbols indicate oseltamivir and etanercept treatment day, respectively. Animals were killed on day 6 p.i. and lung tissue collected for various analyses. Weight loss (A) and clinical scores (B) were monitored until day 6 p.i. Histopathological scores (C) were derived from microscopic examination of the lung histology H&E sections (E-I), where sections were examined using bright field microscope on all fields at 200x magnification. Viral load (D) data was log -transformed. Data are expressed as means ± SEM and were analyzed using two-way ANOVA (A and B) or one-way ANOVA (C and D) with Holm-Sidak's multiple comparisons tests. *, $p < 0.05$; **, $p < 0.01$; and ****, $p < 0.0001$. Broken line in panel D corresponds to the limit of virus detection. Bars in panels E-I correspond to 100 μm. Data shown are from a single experiment.

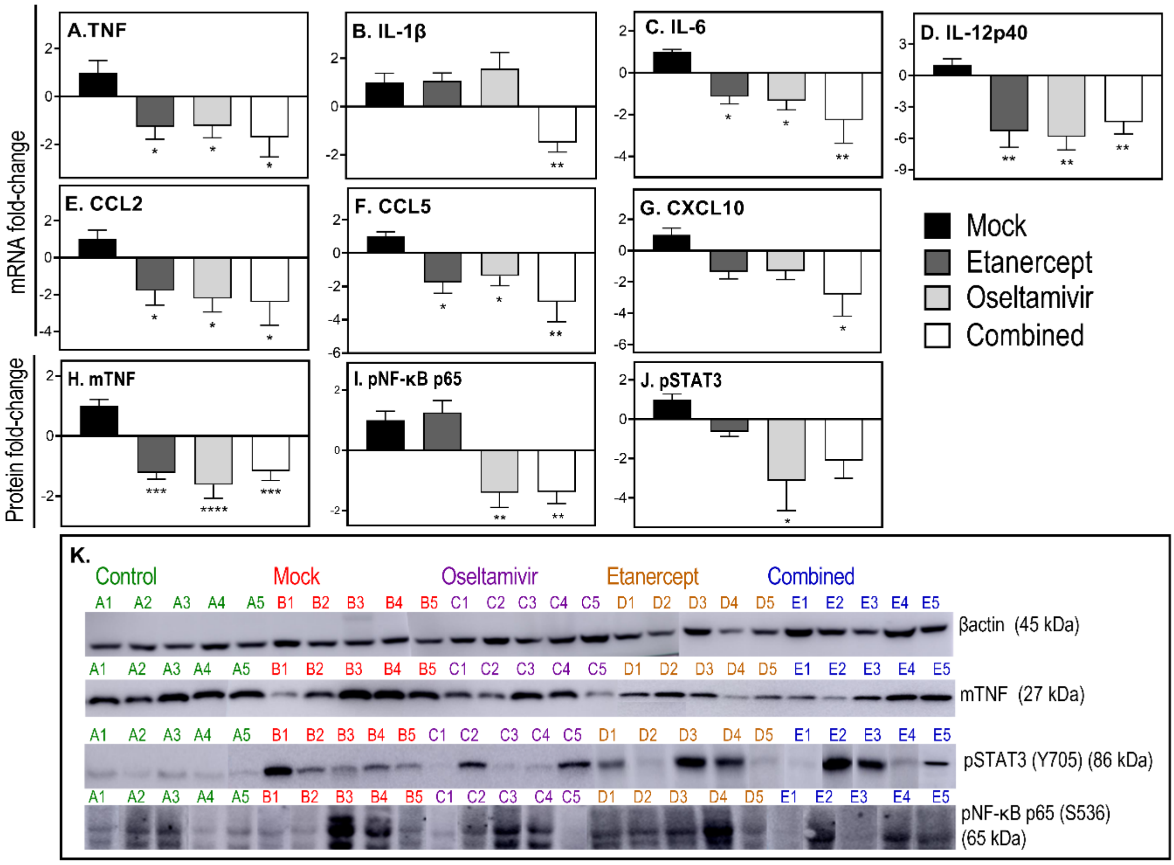


**Fig. S4. Combined treatment with standard dose oseltamivir and one dose etanercept downregulates mRNA transcripts for pro-inflammatory cytokines/chemokines and reduces NF-κB and STAT3 activation.** Lung tissue sections were obtained from mice that were infected and treated as described in Fig. S3. Briefly, age-matched groups of female WT mice were infected with 3000 PFU of IAV i.n. Treatment with etanercept, oseltamivir, or both drugs (combined) were commenced on day 3 p.i. and on days 4 and 5 p.i. only oseltamivir was administered. Animals were killed on day 6 p.i. and lung tissue collected for quantifying mRNA expression levels for specific cytokines and chemokines using qRT-PCR (A-G) and protein levels of mTNF (H), pNF-κB p65 (I), and pSTAT3 (J) were measured using western blot (K) and quantified with the ImageJ software. For (K), samples were run on 3 separate gels, i.e. gel 1, A1-B4; gel 2, B5-D2; gel 3 D3-E5. Data are expressed as mean fold-change relative to the mock treated group ± SEM of 5 replicates. Statistical analyses were performed using one-way ANOVA with Holm-Sidak's multiple comparisons tests. *, $p < 0.05$; **, $p < 0.01$; ***, $p <0.001$ and ****, $p <0.0001$. Data shown are from a single experiment.

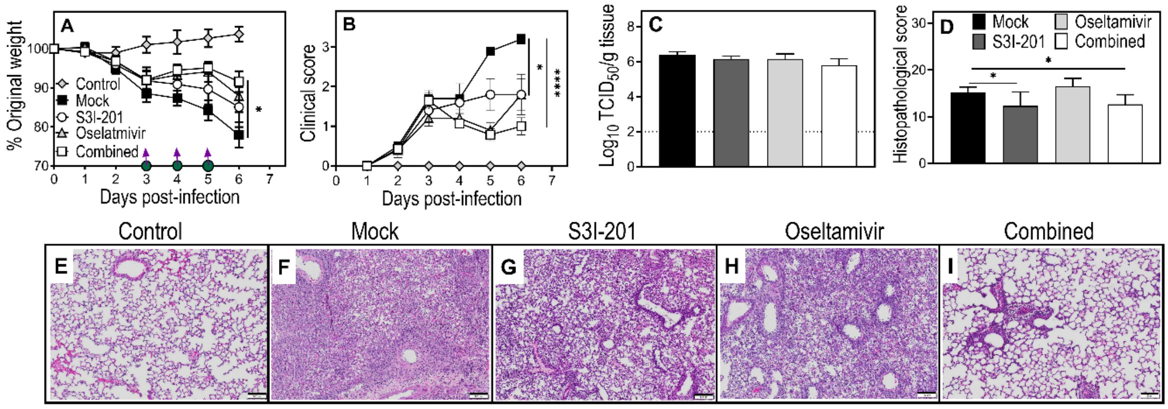


**Fig. S5. Combined treatment with standard dose oseltamivir and S3I-201 reduces weight loss, clinical scores and lung pathology but not viral load.** Age-matched groups of female WT mice were infected with 3000 PFU IAV (n = 5) or given PBS (n = 5) i.n. Animals were treated with S3I-201, oseltamivir (20mg/kg, twice a day), or both drugs (combined) on days 3, 4 and 5 p.i., as indicated in panel A, where a purple arrow and a filled green circle symbols indicate oseltamivir and S3I-201 treatment day, respectively. Animals were killed on day 6 p.i. and lung tissues collected for various analyses. Weight loss (A) and clinical scores (B) were monitored until day 6 p.i. (C) Viral load data were log-transformed. (D) Histopathological scores were derived from microscopic examination of the lung histology H&E sections examined using bright field microscope on all fields at 200x magnification (E-I). Data, expressed as mean ± SEM, were analyzed using two-way (A) and (B) or one-way (C) and (D) ANOVA with Holm-Sidak's multiple comparisons tests. *, $p < 0.05$ and ****, $p < 0.0001$. Broken line in panel C corresponds to the limit of virus detection. Bars in panels E-I correspond to 100 μm. Data shown are from a single experiment.

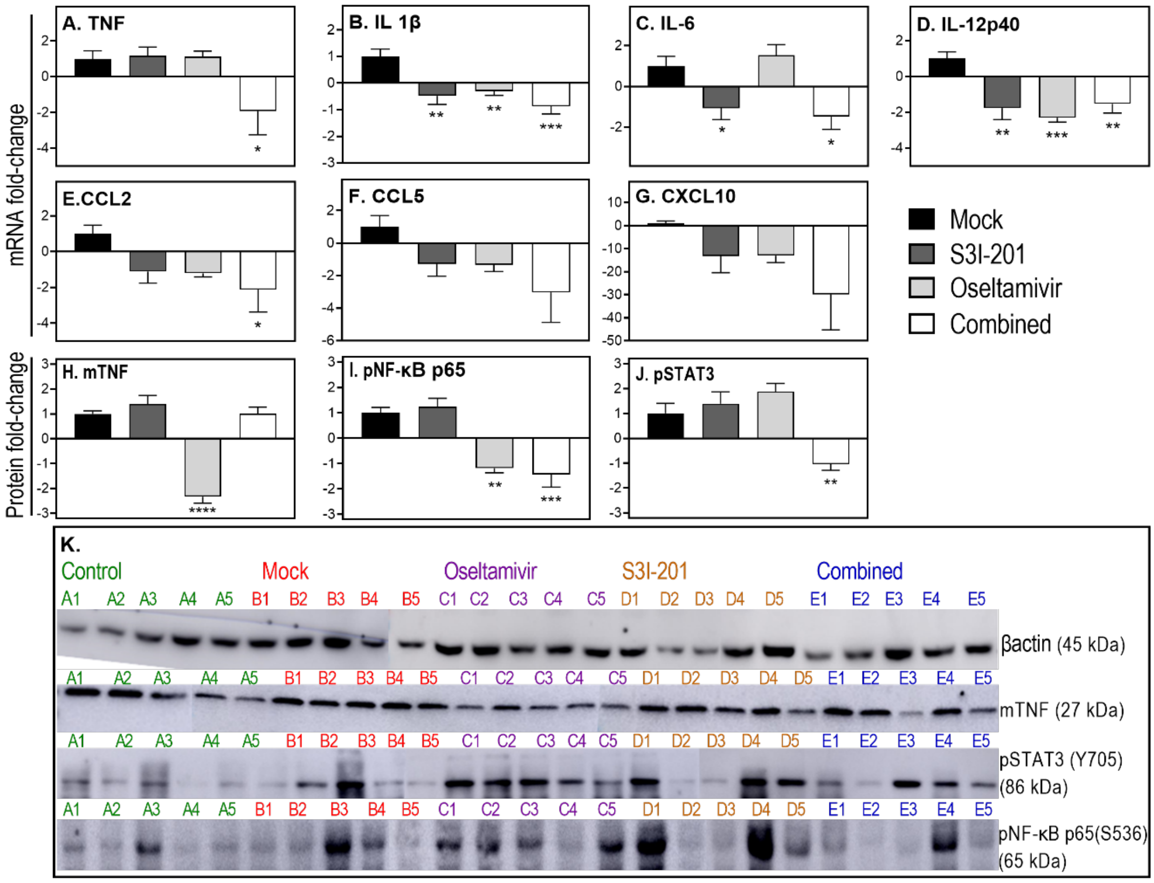


**Fig. S6. Combined treatment with standard dose oseltamivir and S3I-201 reduces mRNA transcripts for pro-inflammatory cytokines/chemokines and activation of NF-κB and STAT3.** Age-matched groups of female WT mice were infected with 3000 PFU IAV (n = 5) or given PBS (n = 5) i.n. Animals were treated with S3I-201, oseltamivir (20 mg/kg, twice a day), or both drugs (combined) on day 3, 4, and 5 p.i. Animals were killed on day 6 p.i. and lung tissues were collected for various analyses. Levels of expression of mRNA transcripts for the indicated cytokines and chemokines were quantified using qRT-PCR (A-G) and protein levels of mTNF (H), pNF-κB p65 (I), and pSTAT3 (J) were measured using western blot (K) and quantified with the ImageJ software. For (K), samples were run on 3 separate gels, i.e. gel 1, A1-B4; gel 2, B5-D4; gel 3 D5-E5 Data are expressed as mean fold-change relative to the mock treated group ± SEM and were analyzed using one-way ANOVA with Holm-Sidak's multiple comparisons tests. *, $p < 0.05$; **, $p < 0.01$; ***, $p < 0.001$ and ****, $p < 0.0001$. Data shown are from a single experiment.

### Supplementary Information References


1. A. L. Balish, J. M. Katz, & A. I. Klimov Influenza: propagation, quantification, and storage. *Current protocols in microbiology* **29**, 15G. 11.11-15G. 11.24 (2013).
2. L. J. Reed & H. Muench A simple method of estimating fifty percent endpoints. *American Journal of Epidemiology* **27**, 493-497 (1938).
3. J. C. Hierholzer & R. A. Killington (1996) 2 - Virus isolation and quantitation. *Virology Methods Manual*, eds Mahy BWJ & Kangro HO (Academic Press, London), pp 25-46.
4. M. J. Tuazon Kels *et al.,* TNF deficiency dysregulates inflammatory cytokine production, leading to lung pathology and death during respiratory poxvirus infection. *Proc Natl Acad Sci U S A* **117**, 15935-15946 (2020).
5. Z. Al Rumaih *et al.,* Poxvirus-encoded TNF receptor homolog dampens inflammation and protects from uncontrolled lung pathology during respiratory infection. *Proceedings of the National Academy of Sciences* **117**, 26885-26894 (2020).
6. K. J. Livak & T. D. Schmittgen Analysis of relative gene expression data using real-time quantitative PCR and the 2(-Delta Delta C(T)) Method. *Methods* **25**, 402-408 (2001).
7. H. Davarinejad Quantifications of Western Blots with ImageJ. Available online: http://www.yorku.ca/yisheng/Internal/Protocols/ImageJ.pdf (accessed on 28/06/2021).
